# Adaptive anti-tumor immunity is orchestrated by a population of CCL5-producing tissue-resident NK cells

**DOI:** 10.1101/2021.05.27.445981

**Authors:** Nicole Kirchhammer, Marcel P Trefny, Marina Natoli, Dominik Brücher, Sheena N Smith, Franziska Werner, Victoria Koch, David Schreiner, Ewelina Bartoszek, Mélanie Buchi, Markus Schmid, Daniel Breu, K Patricia Hartmann, Polina Zaytseva, Daniela S Thommen, Heinz Läubli, Jan P Böttcher, Michal A Stanczak, Abhishek S Kashyap, Andreas Plückthun, Alfred Zippelius

## Abstract

T cell-directed cancer immunotherapy often fails to generate lasting tumor control. Harnessing additional effectors of the immune response against tumors may strengthen the clinical benefit of immunotherapies. Here, we demonstrate that therapeutic targeting of the IFNγ-IL-12 pathway relies on the ability of a population of tissue-resident NK (trNK) cells to orchestrate an anti-tumor microenvironment. Particularly, utilizing an engineered adenoviral platform, we show that paracrine IL-12 enhances functional DC-CD8 T cell interactions to generate adaptive anti-tumor immunity. This effect depends on the abundance of trNK cells and specifically their capacity to produce the cDC1-chemoattractant CCL5. Failure to respond to IL-12 and other IFNγ-inducing therapies such as immune checkpoint blockade in tumors with low trNK cell infiltration could be overcome by intra-tumoral delivery of CCL5. Our findings reveal a novel barrier for T cell-focused therapies and offer mechanistic insights into how T cell-NK cell-DC crosstalk can be enhanced to promote anti-tumor immunity and overcome resistance.

**Significance:** We identified the lack of CCL5-producing, tissue-resident NK (trNK) cells as a barrier to T cell-focused therapies. While IL-12 induces anti-tumoral DC-T cell crosstalk in trNK cell^rich^ tumors, resistance to IL-12 or anti-PD-1 in trNK cell^poor^ tumors can be overcome by the additional delivery of CCL5.

## Introduction

The clinical success of immune checkpoint blockade has initially kept the scientific focus predominantly on factors regulating T cell activity ^1^. It is, however, increasingly acknowledged that a diverse range of immune cells, including components of innate immunity, must function in a coordinated and synergistic manner to successfully achieve immune-mediated tumor rejection ^2–4^.

The interferon γ (IFNγ)-Interleukin 12 (IL-12) axis plays a central role in connecting innate and adaptive cancer immunity ^5^. Mainly produced by dendritic cells (DCs) in the tumor microenvironment, IL-12 stimulates cytotoxicity and cytokine-secretion in T cells and natural killer cells (NK cells) ^6^. In a positive IL-12-IFNγ feedback loop, T and NK cell-derived IFNγ in turn activates and induces IL-12 expression in DCs. Moreover, IFNγ enhances antigen (cross-)presentation by antigen-presenting cells (APCs) thereby further potentiating the cytotoxic activity of CD8 T cells ^5,7^. Consequently, gene expression signatures reflecting cellular components of this axis — NK cells, DCs, and CD8 T cells —, as well as signatures of IFNγ signaling are predictive of improved patient survival in multiple cancer types ^8–12^. Furthermore, it has been shown that IL-12 induction by IFNγ is essential for the efficacy of immune checkpoint blockade ^5^.

IL-12 has been extensively investigated for its use in cancer immunotherapy and has demonstrated remarkable antitumor efficacy in tumor models. However, in early clinical trials, its therapeutic benefit in patients remained limited with severe dose-limiting toxicity ^13,14^. A possible explanation is a lack of targeting to the tumor microenvironment. Most cytokines, including IL-12, act locally in the tumor and nearby lymph nodes in a paracrine or autocrine fashion, rather than systemically ^15^. While multiple approaches using localized IL-12 delivery are currently under investigation, the striking anti-tumor efficacy of local IL-12 therapy observed in mice has yet to be replicated in humans ^16–19^.

We here sought to gain a deeper understanding of how IL-12-mediated tumor control is achieved and whether this knowledge ultimately allows to design improved treatment strategies for efficient tumor control. To this end, we utilized a tumor-targeted adenovirus serotype 5 delivery platform as a tool for intra-tumoral IL-12 immunotherapy (AdV5-IL12) ^20–22^. In tumor models and patient-derived model systems, we demonstrate that the efficacy of IL-12 depends on the intra-tumoral abundance of a population of tissue-resident NK (trNK) cells and their ability to prime the immune microenvironment by producing the chemokine CCL5. In trNK cell^poor^ tumors, resistance to IL-12 can be rescued by induced expression of CCL5, leading to increased infiltration of cDC1s, which then set the positive anti-tumor DC-T cell feedback loop in motion. Similarly, resistance to other IFNγ-mediated treatments such as PD-1 checkpoint blockade can be a result of reduced CCL5 induction due to a lack of trNK cells and can be overcome by treatment with CCL5. Our data highlight the importance of tumor-resident NK cells for the induction of DC-T cell crosstalk and successful cancer immunotherapy.

## Results

### IL-12 immunotherapy prompts NK cells to orchestrate an anti-tumor microenvironment

Even though strategies to maximize IL-12 delivery are of increasing clinical interest, the therapeutic benefit in patients remains moderate. We first sought to identify cell types and key pathways underlying successful clinical outcomes to intra-tumoral IL-12 therapy in melanoma patients (IL-12MEL trial; NCT01502293) ^16^. To this end, we correlated tumor immune signatures derived from pre-treatment tumor biopsies with therapeutic responses. Patients with clinical responses showed higher NK cell and CD8 T cell scores compared to patients with tumor progression, while no correlation was found with scores of other immune cells (Figure 1a and Supplementary Fig. S1a).

**Figure 1:**
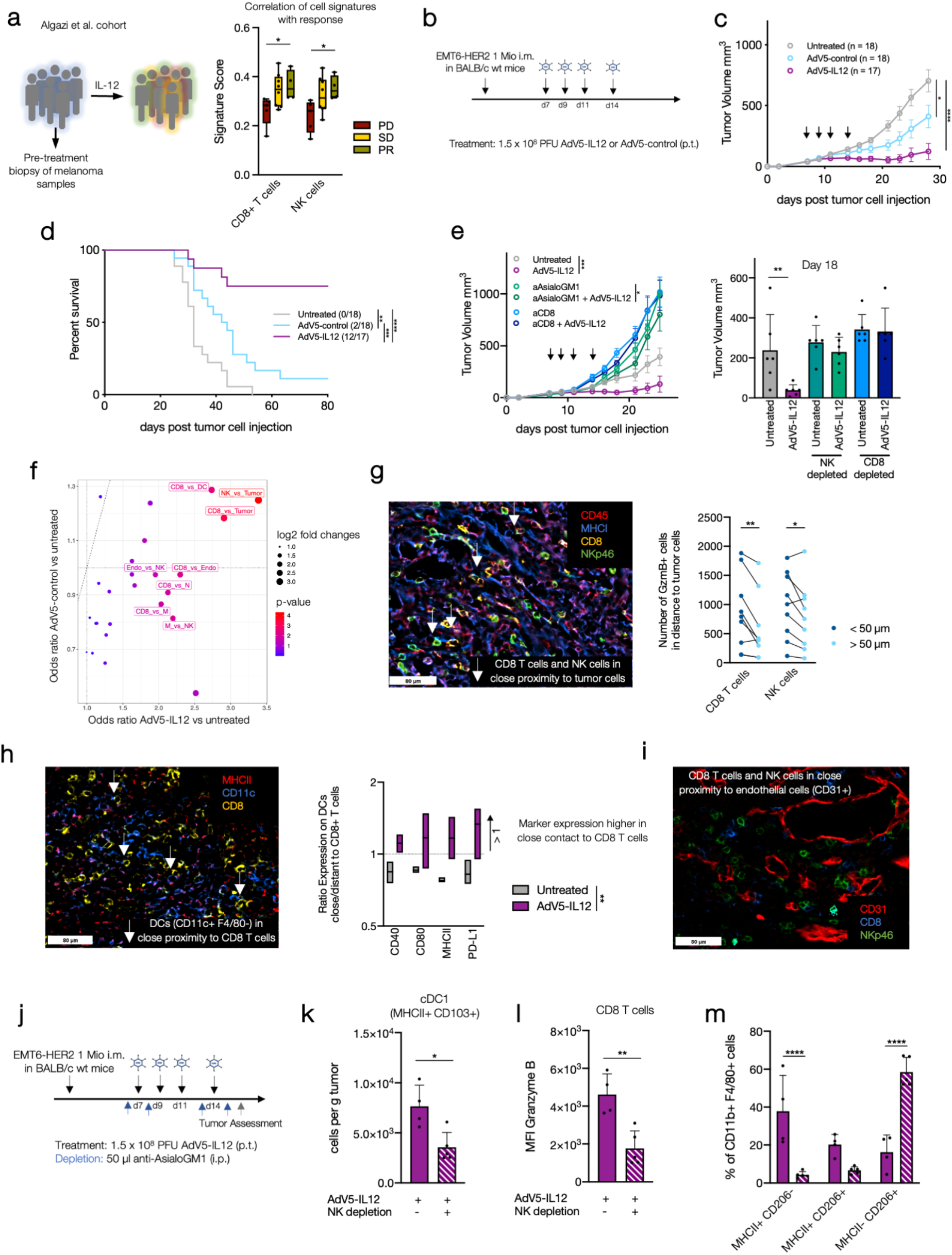
IL-12 immunotherapy prompts NK cells to orchestrate an anti-tumor microenvironment. **a:** Cell signature scores measured by Nanostring in skin tumor biopsies from 19 melanoma patients before intra-tumoral treatment with ImmunoPulse IL-12 were correlated with clinical response (PD: progressive disease, SD: stable disease, PR: partial response) ^16^. **b-d:** Wildtype (WT) mice were engrafted with 1 mio EMT6-HER2 intramammarily (i.m.). From day 7 (tumor size 30–70 mm^3^), mice were treated with 1.5×10^8^ PFU of HER2-targeted and shielded adenoviral vectors (peritumorally, p.t.) encoding for IL-12 or an empty control cassette (AdV5-control) on days 7, 9, 11 and 14 p.t. Tumor growth and Kaplan-Meier survival curves are shown with the number of mice indicated. Black arrows denote days of treatment. e: For depletion studies, mice were injected i.p. with anti-CD8 and anti-AsialoGM1 starting one day before adenoviral treatment. Tumor growth curves after depletion are shown. Black arrows denote days of adenoviral treatment. n = 6 mice. f: Visualization of odds ratios and p values for changes in cell-cell type interactions between experimental conditions focusing on interaction including CD8 T cells and NK cells. g: Representative IF pictures are showing AdV5-IL12 treated tumors (CD45: red, MHCI: blue, CD8: yellow, NKp46: green). White arrows are showing CD8 T cells (CD45+, MHCI+, CD8+) or NK cells (CD45+, MHCI+, NKp46+) neighboring tumor cells (CD45-, MHCI+). Quantification of GzmB+ CD8 T cells and NK cells in close proximity (<50 μm) to tumor cells in comparison to more distant (>50 μm) proximity. Each dot represents the count in one acquired tumor. Treatment conditions were pooled in this analysis. h: Representative IF pictures are showing AdV5-IL12 treated tumors (MHCII: red, CD11c: blue, CD8: yellow). White arrows are showing CD8 T cells (CD45+, CD8+) neighboring DCs (CD45+, CD11c+, F4/80-). Ratio of CD40, CD80, MHCII and PD-L1 expression on the DC cluster in close (<50 μm) or distant (>50 μm) proximity to CD8 T cell cluster comparing untreated and AdV5-IL12 treated tumors are shown. i: Representative IF pictures showing CD8 T cells and NK cells in close proximity to blood vessels in AdV5-IL12 treated tumors (CD31: red, CD8: blue, NKp46: green). f-i: n = 3 mice per group. j: EMT6-HER2-bearing mice were treated with AdV5-IL-12. NK cells were depleted using anti-AsialoGM1 antibody (i.p) as indicated (blue arrow). Tumors were isolated and single cell suspensions of tumors digest were analyzed using flow cytometry on day 16. k: Intra-tumoral cDC1s (CD11c+, F4/80-, Ly-6G-, MHCII+, CD103+, CD11b^low^) were quantified after AdV5-IL12 treatment +/− NK depletion. l: Mean fluorescence intensity (MFI) of granzyme B on CD8 T cells (CD3+, CD8+, NKp46-, CD19-, Ly-6G-). m: Polarization of macrophages (CD11b+ F4/80+) after AdV5-IL12 treatment comparing NK depletion versus non-depleted. k-m: n = 4-5 mice per group. *p < 0.05, **p < 0.01, ***p < 0.001, ****p < 0.0001. Error bar values represent SD or SEM (tumor growth curves). For comparisons between three or more groups, one-way ANOVA with multiple comparisons was used. For survival analysis, p values were computed using the Log Rank test. Two-way ANOVA was used to compare tumor growth curves. See also Supplementary Fig. S1–4.

To define a model system for local IL-12 therapy that reflects these clinical findings, we utilized a non-replicative, shielded, and re-targeted adenovirus serotype 5 vector previously established in our laboratory ^20–22^. For this study, HER2 was used as a model antigen to target the HER2-overexpressing syngeneic tumor cell lines B16-HER2 and EMT6-HER2. Due to the high abundance of pre-existing antibodies against AdV5, we made use of a shield based on a hexon-binding humanized single-chain variable fragment (scFv), fully covering the virion ^22^. To confirm tumor-specific expression of our payload, luciferase-encoding virus (AdV5-Luciferase) was peritumorally injected into B16-HER2 and EMT6-HER2 bearing mice (Supplementary Fig. S1b). In both tumor models, the payload was exclusively expressed in the tumor for up to 10 days with a peak expression on day 1 (Supplementary Fig. S1c–e).

To evaluate the efficacy of AdV5 encoding IL-12 (AdV5-IL12), we peritumorally treated mice bearing orthotopic EMT6-HER2 tumors with four injections of 1.5×10^8^ PFU retargeted and shielded AdV5-IL12. Empty virus (AdV5-control) served as a control (Figure 1b). Treatment with AdV5-IL12 resulted in strong inhibition of tumor growth, enhanced survival, and complete tumor regression in 70% of treated mice, while AdV5-control showed only moderate effects (Figure 1c–d). No IL-12 was detected in the serum, confirming the specificity of our tumor-localized therapy (Supplementary Fig. S1f). Retargeted and shielded AdV5-IL12 showed increased efficacy compared to naked and retargeted vectors (Supplementary Fig. S1g).

As patients with localized IL-12 therapy show clinical responses even in non-treated lesions, we assessed the systemic effects of AdV5-IL12 in our mouse tumor model. To this end, we injected HER2-negative EMT6 wt cells into the contralateral flanks of the EMT6-HER2 tumors (Supplementary Fig. S1h). We observed reduced tumor growth of the contralateral tumors and complete regression in 50% of the AdV5-IL12 treated animals. This indicates potent systemic immune effects upon local administration of AdV5-IL12.

AdV5-IL12 induced the formation of protective anti-tumor immunological memory as mice that survived primary EMT6-HER2 engraftment following AdV5-IL12 treatment remained tumor-free after later re-challenge with the same cell line (Supplementary Fig. S1i). Tumors from mice simultaneously inoculated with EMT6 wt cells on the lateral flank were equally rejected suggesting broad memory formation against shared antigens expressed by EMT6 cells (Supplementary Fig. S1i).

To dissect the role of defined immune cell populations in mediating the therapeutic effect of AdV5-IL12, we performed antibody-mediated depletion studies (Figure 1e). In agreement with the immune signature analysis from the IL-12MEL trial (Figure 1a), AdV5-IL12 required both CD8 T cells and NK cells for therapeutic efficacy. As IL-12 is a known driver of IFNγ production in both those cell types and the clinical response correlates with a defined IFNγ score ^16^, we assessed the contribution of IFNγ to the activity of AdV5-IL12. IFNγ-neutralized mice failed to control EMT6-HER2 tumors upon treatment with AdV5-IL12 (Supplementary Fig. S1j). In line with these findings, AdV5-IL12 increased the capacity of CD8 T cells and NK cells to proliferate and to exert effector functions as assessed by flow cytometry (Supplementary Fig. S2 and S3). In conclusion, mirroring the clinical situation, AdV5-IL12 treatment in EMT6-HER2 tumors requires robust NK and CD8 T cell responses which depend on IFNγ.

We next assessed the capability of CD8 and NK cells to directly interact and attack cancer cells using a highly multiplexed cytometric imaging approach, termed co-detection by indexing (CODEX) ^23^. AdV5-IL12 led to a pronounced accumulation of CD45+ immune cells (Supplementary Fig. S4a–b). Spatial proximity (defined as a distance of <50 μm) of NK and CD8 T cells with tumor cells was specifically increased upon AdV5-IL12 treatment (Figure 1f and Supplementary Fig. S4c). To confirm functional interactions and tumor lysis by NK and CD8 T cells, we analyzed the expression of the effector marker granzyme B in cells with spatial proximity to tumor cells (CD45-CD31-). In both NK and CD8 T cells, the number of granzyme B+ cells was increased in close proximity to tumor cells (Figure 1g). In addition, the spatial proximity of CD8 T cells with DCs was specifically enhanced after AdV5-IL12 treatment (Figure 1f). Moreover, the co-stimulatory molecules CD40, CD80 and MHCII, as well as PD-L1 on DCs in close proximity to CD8 T cells were specifically increased after AdV5-IL12 treatment, indicating an enhanced functional interaction between DCs and CD8 T cells induced by IL-12 (Figure 1h). We also noticed close proximity of CD8 and NK to endothelial cells after AdV5-IL12 treatment (Figure 1f and i) which may indicate induced trafficking of these cells by IL-12.

To investigate IL-12 induced immune cell recruitment to the tumor, we blocked lymphocyte recirculation with the trafficking inhibitor FTY720 ^24^. FTY720 treatment prior to tumor inoculation fully abrogated tumor control, while blocking trafficking during AdV5-IL12 treatment allowed initial tumor control, but did not result in complete tumor regression (Supplementary Fig. S4d). This suggests that efficacious IL-12 responses require both pre-existing and actively recruited tumor infiltrating lymphocytes.

In addition to direct cytotoxicity against tumor cells, NK cells were shown to drive immune cell infiltration and intrinsic inflammation within tumors ^2,9^. To dissect the impact of NK cells on IL-12-induced recruitment, we depleted NK cells throughout the AdV5-IL12 treatment and assessed fate and infiltration of tumor-infiltrating immune cells (Figure 1j and Supplementary Fig. S3). NK cell depletion led to a reduced number of MHCII+ CD103+ type I conventional dendritic cells (cDC1s, CD11c+ F4/80-; Figure 1k). Furthermore, NK depletion was associated with reduced granzyme B expression in CD8 T cells, suggesting an important role of NK cells in facilitating optimal anti-tumor CD8 T cells responses (Figure 1l). While the number of macrophages (CD11b+ F4/80+) in the tumor was unchanged, depletion of NK cells skewed the polarization towards an immunosuppressive M2 (CD206+ MHCII-) phenotype (Figure 1m). Thus, we concluded that during AdV5-IL12 treatment, NK cells orchestrate the recruitment and priming of essential cell subsets including cDC1s, and activated CD8 T cells to enhance tumor killing.

### CCL5, mainly produced by trNK cells, is required for IL-12 mediated tumor rejection

To identify clinically relevant chemokines which may guide immune cell recruitment by NK cells after IL-12 therapy, we correlated NK cell scores with the expression of chemokines and their receptors in patients responding to IL-12 therapy (Figure 2a). We identified *CCL5* and its receptor *CCR5* strongly correlated with the NK score (Figure 2b). Accordingly, *CCL5* was significantly upregulated in patients with clinical responses (Figure 2c). These results suggest that NK cells, likely through CCL5, are important in facilitating clinical responses to IL-12.

**Figure 2:**
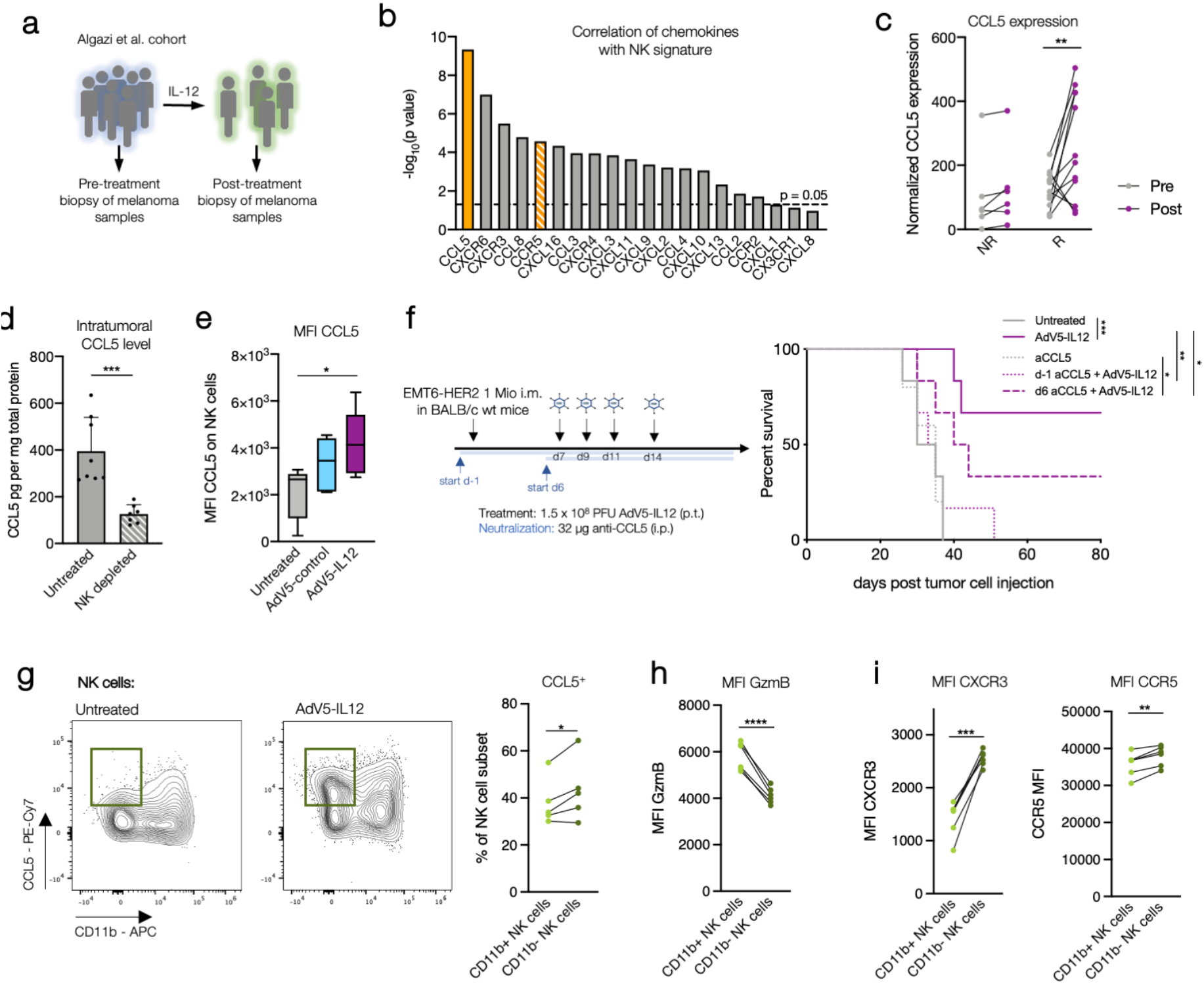
CCL5, mainly produced by trNK cells, is required for IL-12 mediated tumor rejection. **a-b:** −log10(p value) of chemokines and their receptors which are correlating with NK signature of skin tumor biopsies from melanoma patients responding to intra-tumoral treatment with ImmunoPulse IL-12 are shown. c: Normalized counts of *CCL5* expression of patients with no-response (NR: PD) versus response (R: SD + PR) before and after intra-tumoral treatment with ImmunoPulse IL-12 determined by NanoString. d: NK cells were depleted using anti-AsialoGM1 antibody (i.p) every 3-4 days. On day 7 post EMT6-HER2 inoculation (tumor size 30–70 mm^3^), tumors were isolated and lysed. CCL5 expression was determined by ELISA and normalized to total protein measured by BCA. n= 6 per condition. 2-tailed Student’s t test was used. e: Mice were treated with AdV5-IL12 or AdV5-control on day 7, 9 and 11 post EMT6-HER2 inoculation. On day 12 tumors were isolated and CCL5 expression (MFI) was analyzed on NK cells. n= 6 mice per condition. f: EMT6-HER2-engrafted mice were treated with AdV5-IL12 following the indicated schedule. Starting one day prior tumor inoculation or one day prior adenoviral therapy, CCL5 was neutralized using antibodies every 3-4 days. Kaplan-Meier survival curves are shown. g-i: Mice were treated with AdV5-IL12 or AdV5-control on day 7, 9 and 11 post EMT6-HER2 inoculation. On day 12 tumors were isolated and CCL5, GzmB, CXCR3 and CCR5 expression was analyzed on NK cell subsets. Percentage of CCL5 positive NK cells, as well as MFI of GzmB, CXCR3 and CCR5 were quantified after AdV5-IL12 treatment. Paired 2-tailed Student’s t test was used. *p < 0.05, **p < 0.01, ***p < 0.001, ****p < 0.0001. Error bar values represent SD or SEM (tumor growth curves). For comparisons between three or more groups, one-way ANOVA with multiple comparisons was used. For survival analysis, p values were computed using the Log Rank test. Two-way ANOVA was used to compare tumor growth curves. See also Supplementary Fig. S4.

We therefore investigated whether NK cell-mediated CCL5 contributes to the efficacy of AdV5-IL12 in EMT6-HER2 tumors. We first showed that CCL5 concentrations were dramatically reduced in tumor lysates after NK cell depletion, which confirms NK cells as the main source of CCL5 in this model (Figure 2d). In line with the observed CCL5 upregulation in responding patients (Figure 2c), AdV5-IL12 treatment of EMT6-HER2 tumors induced CCL5 production in NK cells (Figure 2e). Neutralizing CCL5 before tumor inoculation fully abrogated IL-12 efficacy, while CCL5 neutralization after tumor inoculation but before AdV5-IL12 treatment partially inhibited efficacy (Figure 2f). Therefore, we concluded that CCL5 produced by NK cells plays a dual role in the response to IL-12 delivery. While CCL5 at the steady-state permits responsiveness to AdV5-IL12 treatment, further induction of CCL5 by AdV5-IL12 treatment may attract immune cells to improve anti-tumor immunity.

To analyze the contribution of CCL5 to NK cell-mediated tumor rejection upon AdV5-IL12 treatment, NK-depleted tumor-bearing mice were concomitantly treated with AdV5-IL12 and an AdV5 encoding CCL5 (AdV5-CCL5). While depletion of NK cells abrogated the efficacy of AdV5-IL12 (Fig 1c), the combination of AdV5-CCL5 with AdV5-IL12 partly rescued the efficacy (Supplementary Fig. S4e).

Recent studies have revealed that NK cells are highly heterogeneous with different immune functions ^25–27^. To identify which NK cell subset produced CCL5, we analyzed CCL5 levels after AdV5-IL12 treatment in subpopulations of NK cells reflecting different maturation stages, defined by CD11b expression ^28^. We observed a higher proportion of non-mature (CD11b-) NK cells producing CCL5 compared to mature (CD11b+) cells (Figure 2g). CD11b-NK cells showed a lower expression of cytotoxic mediators (granzyme B, Figure 2h) and a higher expression of chemokine receptors associated with tissue entry and retention (CXCR3 and CCR5, Figure 2i) ^29^. These findings suggest that responsiveness to IL-12 depends on CCL5-producing NK cells which exhibit features associated with tissue-residency with lower cytotoxic profile (in the following defined as tissue-resident NK cells).

### AdV5-CCL5 overcomes AdV5-IL12 resistance in trNK cell poor tumors

We thus hypothesized that tumors with a lower proportion of tissue-resident (trNK) cells would largely be resistant to AdV5-IL12. To test this, we determined the impact of NK cell depletion on the efficacy of AdV5-IL12 in B16-HER2 tumors, a tumor model with a lower proportion of trNK cells but a similar total number of NK cells, compared to the EMT6-HER2 model (Figure 3a–b). In line with our findings that trNK cells are the major source of CCL5, CCL5 levels were lower in B16-HER2 tumors compared to EMT6-HER2 tumors (Figure 3c). Unlike in EMT6-HER2 tumors, NK cell depletion in the B16-HER2 model did not increase tumor growth of untreated or AdV5-IL12 treated tumors suggesting that mainly trNK cells but not mature conventional NK (cNK) cells determine IL-12 responsiveness (Figure 1e and 3d).

**Figure 3:**
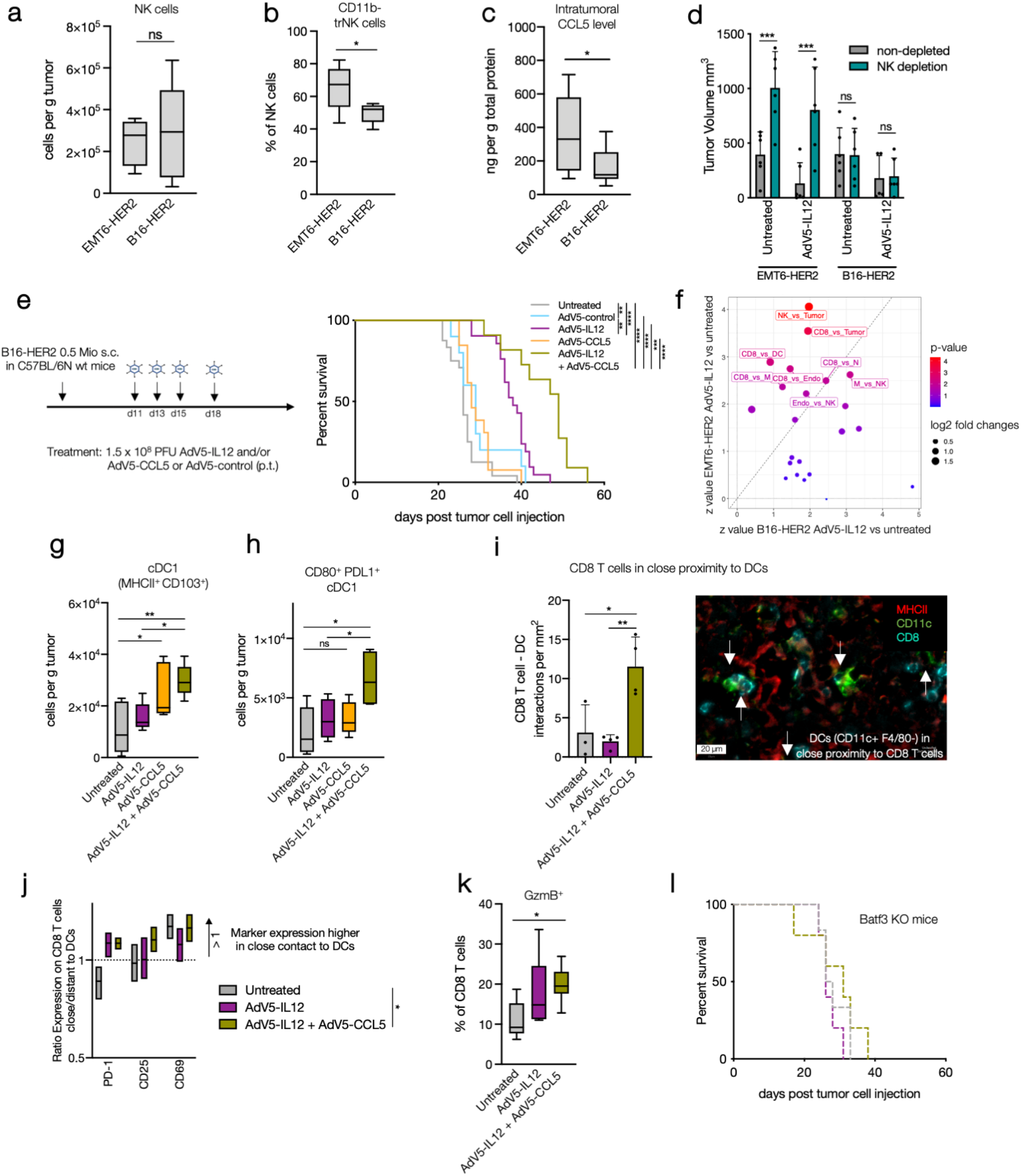
AdV5-CCL5 overcomes AdV5-IL12 resistance in in trNK cell poor tumors. **a:** Amount of tumor infiltration NK cells per g tumor was determined in untreated EMT6-HER2 and B16-HER2 tumors. **b:** Amount of trNK cells per g tumor was determined in untreated EMT6-HER2 and B16-HER2 tumors. **c:** Intra-tumoral CCL5 concentration was determined by ELISA and normalized to total protein in EMT6-HER2 and B16-HER2 tumor lysates. **d:** Wildtype (WT) mice were engrafted with 1 mio EMT6-HER2 (i.m.) or 0.5 mio B16-HER2 (s.c.). Mice were treated with AdV5-IL12 on day 7, 9, 11 and 14 or day 11, 13, 15 and 18 (tumor size 30–70 mm^3^), respectively. NK cells were depleted using anti-AsialoGM1 and anti-NK1.1 antibody every 4-5 days starting one day prior adenoviral therapy. Tumor volume on day 25 post tumor inoculation is shown. e: Mice were treated with AdV5-IL12 and AdV5-CCL5 on day 11, 13, 15 and 18 (tumor size 30–70 mm^3^) after B16-HER2 inoculation as indicated: Tumor growth and Kaplan-Meier survival curves are shown (n > 15 mice per condition). **f:** Mice were engrafted with 1 mio EMT6-HER2 (i.m.) or 0.5 Mio B16-HER2 (s.c.). Starting from day 7 or 11 (tumor size 30–70 mm^3^), mice were treated with 1.5×10^8^ PFU of HER2-targeted and shielded adenoviral vectors (p.t.) encoding for IL-12 on day 7/11, 9/13 and 11/15. On day 12/16 post inoculation, tumors were isolated, embedded in OCT and analyzed by multiparameter immuno-fluorescence microscopy. Visualization of odds ratios and p values for changes in cell-cell type interactions between EMT6-HER2 versus B16-HER2 focusing on interaction including CD8 T cells and NK cells. **g-k:** Mice were engrafted with 0.5 mio B16-HER2 (s.c.). Mice were treated with AdV5-IL12 and/or AdV5-CCL5 on day 11, 13 and 15 (tumor size 30–70 mm^3^). On day 16 post inoculation, tumors were isolated and single cell suspensions were analyzed by flow cytometry or embedded in OCT and analyzed by multiparameter immuno-fluorescence microscopy. **g-h:** Number of cDC1s (CD11c+, F4/80-, Ly-6G-, MHCII+, CD103+, CD11b^low^) and PD-L1 and CD80 expressing cDC1s, respectively. i: Interaction count per mm^2^ of CD8 T cells in close proximity to DCs. Representative IF pictures are showing AdV5-IL12 + AdV5-CCL5 treated tumors (MHCII: red, CD11c: green, CD8: blue). White arrows are showing CD8 T cells (CD45+, CD8+) neighboring DCs (CD45+, CD11c+, F4/80-). j: Ratio of PD-1, CD25 and CD69 expression on the CD8 T cell cluster in close (<50 μm) or distant (>50 μm) proximity to DC cluster. k: Proportion of granzyme B+ of CD8 T cells (CD3+, CD4-, NKp46-CD19-). l: Batf3 knockout mice (lacking cDC1s) were engrafted with 0.5 mio B16-HER2 (s.c.). Mice were treated with AdV5-IL12 and/or AdV5-CCL5. Tumor growth and Kaplan-Meier survival curves are shown. #x002A;p < 0.05, **p < 0.01, ***p < 0.001, ****p < 0.0001. Error bar values represent SEM. For survival analysis, p values were computed using the Log Rank test. Two-way ANOVA was used to compare tumor growth curves. See also Figure Supplementary Fig. S5.

We then asked if therapeutic CCL5 delivery using AdV5-CCL5 may compensate for the lack of trNK cells when combined with AdV5-IL12 in B16-HER2 bearing mice (Figure 3e). Indeed, the combination further delayed tumor growth and increased survival compared to AdV5-IL12 alone. FTY720 diminished the benefit of AdV5-CCL5/AdV5-IL12, suggesting that CCL5 mainly promotes immune cell recruitment to the tumor (Supplementary Fig. S5a).

To define factors that could explain the lack of efficacy of IL-12 in the trNK cell^poor^ tumors, we compared which cell interactions were induced after IL-12 therapy in the B16-HER2 compared to the EMT6-HER2 mouse tumor model (Figure 3f and Supplementary Fig. S5b–c). We observed two major differences in the changes to the interactome in B16-HER2 tumors. We noticed lower induction of NK-tumor and CD8 T-tumor, and lower induction of DC-CD8 T cell interactions in the trNK cell^poor^ tumor microenvironment. Thus, trNK cell^poor^ B16-HER2 tumors are characterized by a reduced tumor attack of effector cells, and, more importantly, reduced induction of DC-CD8 T cell crosstalk by IL-12. Next, we asked whether CCL5 supplementation in trNK cell^poor^ B16-HER2 tumors can potentially induce DC-CD8 T cell crosstalk and therefore promote anti-tumor T cell immunity. Utilizing multicolor flow cytometry, we were able to detect increased numbers of cDC1s when tumors were treated with either AdV5-CCL5 alone or in combination with AdV5-IL12 (Figure 3g). Accordingly, we found an increased number of cDC1s expressing CD80 and PD-L1 in the AdV5-CCL5-AdV5-IL12 combination (Figure 3h and Supplementary Fig. S5b-c), which indeed led to more interactions between DCs and CD8 T cells (Figure 3i). These interactions resulted in more activated CD8 T cells in proximity to DCs (Figure 3j) leading to an increased proportion of GzmB+ CD8 T cells. In addition, we could observe an increased number of interactions between DCs and tumor cells (Supplementary Fig. S5d). To further understand the contribution of DCs to the therapeutic efficacy of the AdV5-IL12-AdV5-CCL5 combination, we used Batf3 KO mice, lacking cDC1s ^30^. Strikingly, the beneficial effect of AdV5-IL12 +/− AdV5-CCL5 on tumor control and survival was lost in Batf3 KO mice (Figure 3l). Taken together, these data suggest that cDC1s attracted by CCL5 are essential for IL-12-mediated therapeutic benefit by enhancing T cell-mediated immunity; CCL5 can be provided by endogenous trNK cells or therapeutically supplemented in a trNK cell^poor^ environment.

To show the potential of our adenoviral vector as a platform for combinatorial approaches, we then designed adenoviral vectors expressing both, IL-12 and CCL5 (Supplementary Fig. S5e). To avoid deletion of IL-12 or CCL5 by homologous recombination, we encoded both transgenes under the control of orthogonal promotors, CMV or SV40. We were able to demonstrate similar therapeutic efficacy by expressing both payloads within one viral vector, independent of the choice of the promotor, although we thereby decreased the total viral load per injection (Supplementary Fig. S5f–g).

### AdV5-huIL-12 induces CCL5 expression in trNK cells in patient-derived tumor cultures

To test whether a human IL-12-encoding HER2-targeted and shielded adenoviral vector (AdV5-huIL12) can induce similar anti-tumor effects in a human *ex vivo* system, we co-cultured primary tumor suspensions from non-small cell lung cancer (NSCLC) patients (Figure 4a) with a HER2-expressing ovarian cancer cell line (OVCAR3), which was transduced with AdV5-huIL12 (Figure 4a–e). This experimental system has been shown to activate primary human lymphocyte subsets and trigger human cancer cell killing in response to PD-1/PD-L1 blockade ^31^. Within patient-derived tumor-infiltrating lymphocytes (TILs) and OVCAR3 co-cultures, AdV5-huIL12 significantly reduced OVCAR3 viability due to enhanced tumor cell killing (Figure 4b). This effect was accompanied by an increase in IFNγ secretion (Figure 4c) and induction of IFNγ-expressing patient-derived CD8 T cells and NK cells (Figure 4d). In line with our murine data, CCL5 was upregulated in NK cells by IL-12 (Figure 4e).

**Figure 4:**
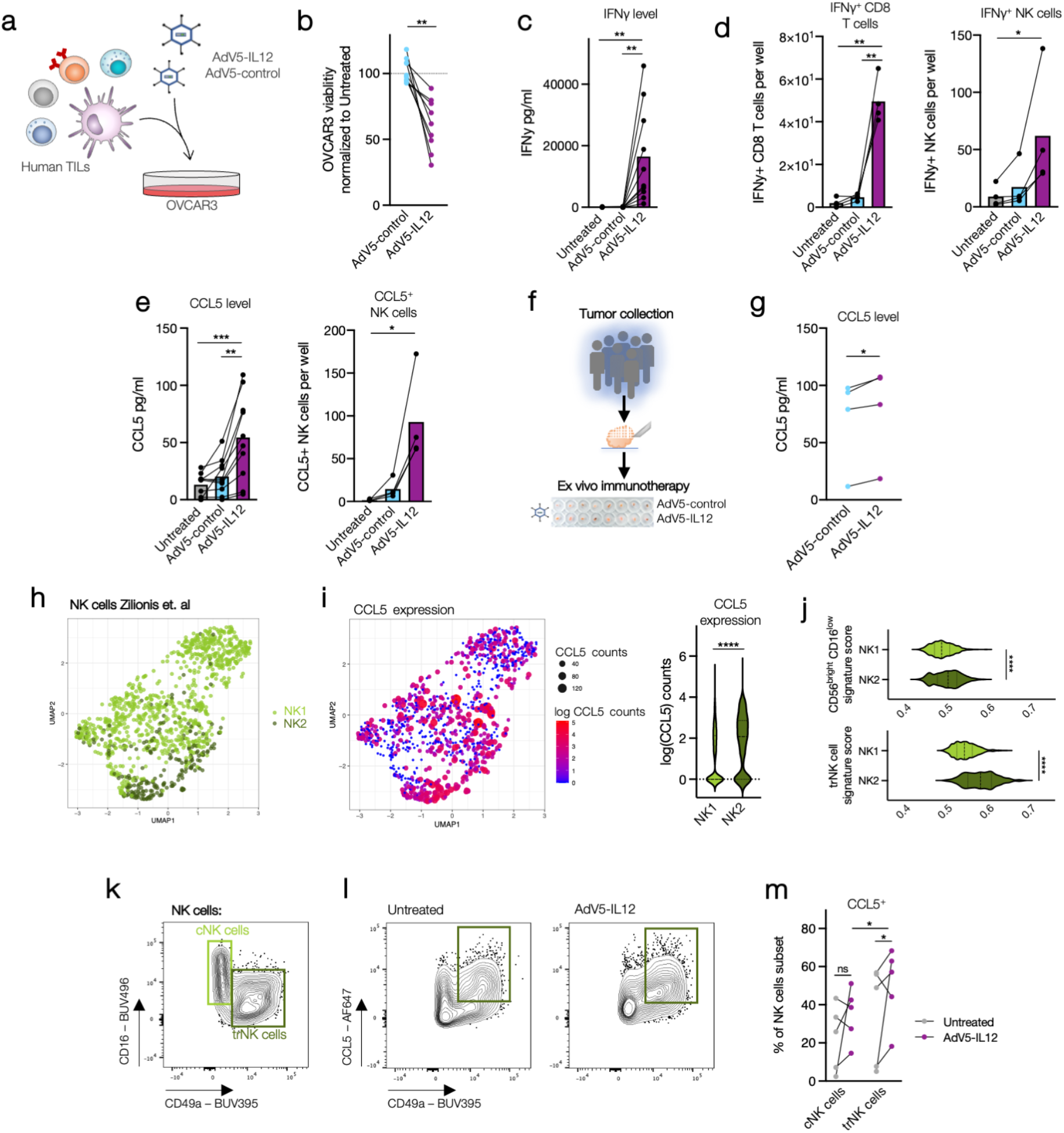
AdV5-huIL-12 induces CCL5 expression in trNK cells in patient-derived tumor cultures. **a-e:** Tumor digests of NSCLC patients were co-cultured with OVCAR-3 cells which were transduced with HER2-targeted AdV5 encoding human IL-12 or control virus. **b:** Quantification of OVCAR-3 viability normalized to untreated co-cultures after 96 h. **c:** IFNγ expression was determined in supernatants after 5d. **d:** Cell count of IFNγ+ CD8 T cells (CD3+, CD56-) and NK cells (CD56+, CD3-) per well after 96 h. **e:** Cell count of CCL5+ NK cells (CD56+, CD3-) after 96 h and CCL5 expression in supernatants after 5d. **f-g:** HER2+ ovarian cancer samples were dissected into tumor fragments and cultivated embedded in matrigel. Tumor fragments were treated with HER2-targeted AdV5 encoding human IL-12 for 48 h (8-12 fragments per condition). **g:** CCL5 concentration in supernatant was analyzed by ELISA. **h:** UMAP projection of scRNASeq data of tumor-infiltrating NK cells of NSCLC patients is shown. **i:** *CCL5* expression in NK1 and NK2 subpopulations was quantified. **j:** CD56^bright^CD16^low^ and trNK cell signature scores of NK1 and NK2 were determined. **k-m:** Tumor digests of NSCLC patients were co-cultured with OVCAR-3 cells which were transduced with HER2-targeted AdV5 encoding human IL-12. **k:** NK cell subsets were defined by CD16 and CD49a expression. **l:** CCL5 producing NK subsets were visualized after AdV5-IL12 treatment compared to untreated. **m:** Percentage of CCL5-positive NK cell subsets were quantified. *p < 0.05, **p < 0.01, ***p < 0.001, ****p < 0.0001. Error bar values represent SD. Paired 2-tailed Student’s t test was used. For comparisons between three or more groups, one-way ANOVA with multiple comparisons was used. See also Figure Supplementary Fig. S6 and S7.

Next, we assessed the activity of AdV5-huIL12 using a human patient-derived tumor fragment model that preserves the tumor microenvironment and architecture but enables *ex vivo* perturbation by checkpoint blockade ^32^. Tumor fragments of HER2-expressing ovarian cancer samples embedded in Matrigel (Figure 4f and Supplementary Fig. S6a) were transduced with AdV5-huIL12. In all four tested patients, we noticed increased staining for IFNγ in CD8 T cells and CCL5 in NK cells (Supplementary Fig. S6b), the latter confirmed in the supernatant by ELISA (Figure 4g).

The identification of trNK cells as the main producers of CCL5 in the murine tumor models above prompted us to characterize which human tumor-infiltrating NK cells are producing CCL5. We therefore investigated the heterogeneity of tumor-infiltrating NK cells in NSCLC patients using a published single-cell RNA sequencing (scRNASeq) data set ^33^. We identified two main NK cell subsets with distinct gene expression profiles with the NK2 subcluster showing a higher expression of *CCL5* compared to the NK1 subcluster (Figure 4h-i and Supplementary Fig. S7a–b). Recently, NK core signature genes associated with cytokine-producing phenotype (CD56^bright^ CD16^-^) and tissue-residency have been described ^34,35^. We noticed a significant enrichment of CD56^bright^ CD16^-^ and tissue-residency genes in the CCL5-producing NK2 subcluster (Figure 4j). Core tissue-residency signature genes include upregulation of the integrin *ITGA1* (CD49a), *ITGAE* (CD103), *CD69*, and *ENTPD1* (CD39) which distinguishes NK2 from the highly *FCGR3A* (CD16)-expressing NK1 subtype. To investigate whether IL-12 induces CCL5 production in NK cells with tissue-residency characteristics (CD49a^+^ CD16^-^), we characterized human tumor-infiltrating NK cells producing CCL5 in our co-culturing system (Figure 4a and Supplementary Fig. S7). In line with our findings from mouse tumor models, CD49a^+^ CD16^-^ trNK cells were producing significantly higher levels of CCL5 in response to AdV5-huIL12 compared to CD16^+^ CD56^dim^ conventional NK (cNK) cells (Figure 4m). Collectively, these experiments recapitulate our finding that IL-12 can directly stimulate the antitumor activity of tumor-infiltrating T cells and induce CCL5 upregulation in trNK cells in primary human tumors.

### CCL5 expression of trNK cells enhances the efficacy of PD-1 blockade

It has been demonstrated that IL-12-producing DCs, activated by IFNγ-secreting CD8 T cells, are critical for successful responses to anti-PD-1 treatments ^5^. This led us to finally investigate whether the DC attractant CCL5 is associated with efficient tumor responses to anti-PD-1 therapy.

In melanoma patients undergoing nivolumab treatment ^36^, we compared CCL5-induction during treatment between responding (R) and non-responding (NR) patients (Figure 5a). While *CCL5* upregulation was associated with clinical responses (Figure 5b), we noticed a positive correlation between the NK2 signature and *CCL5* expression (Figure 5c) ^9^. We also found a positive correlation between the levels of *CCL5* and cDC1s in treated melanoma patients (Figure 5d). These findings suggest that CCL5 may be produced by intra-tumoral trNK cells in response to PD-1 blockade and subsequently might promote cDC1 recruitment, which is in accordance with our observation in tumor models after IL-12 treatment.

**Figure 5:**
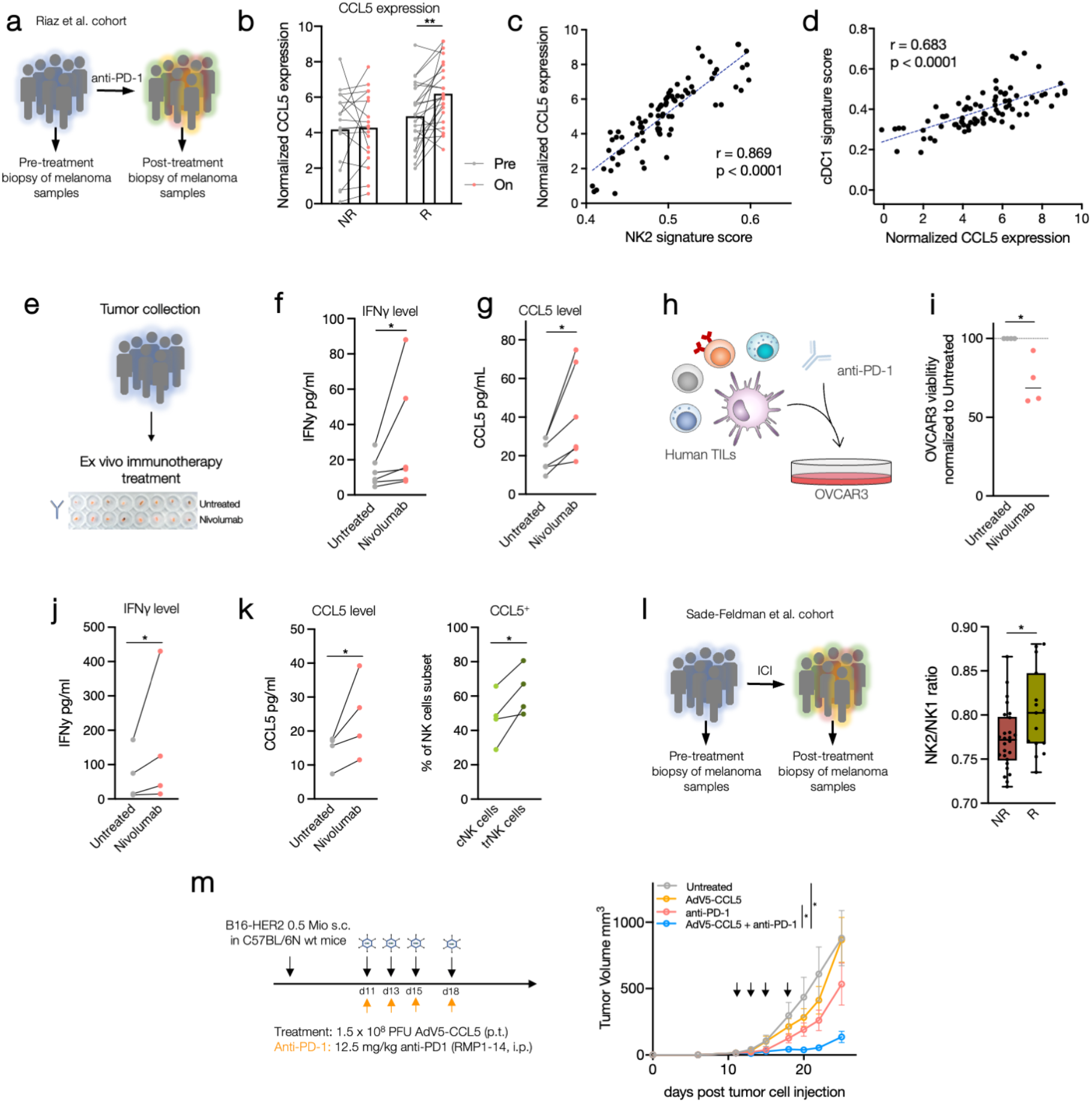
CCL5 expression of trNK cells enhances the efficacy of PD-1 blockade. **a-d:** CCL5 expression in tumor biopsies of melanoma patients (n= 42) was analyzed pre and on nivolumab treatment ^36^. **b:** Normalized expression of *CCL5* pre and on nivolumab treatment between patients with no response (NR) or response (R = SD+PR+CR) was quantified. **c:** Correlation between *CCL5* expression and NK2 signature score or **d:** cDC1 signature score are shown. **e-g:** Cancer samples were dissected into tumor fragments and cultivated embedded in Matrigel. Tumor fragments were treated with nivolumab for 48 h (6-8 fragments per condition). IFNγ and CCL5 were determined in the supernatant. n = 6 tumor samples. **h-k:** Tumor digests of NSCLC patients treated with nivolumab were co-cultured with OVCAR-3 cells for 48h. **i:** OVCAR3 viability was normalized to untreated control group. **j:** IFNγ and **k:** CCL5 were determined in the supernatant. Percentage of CCL5 positive NK cells was quantified after nivolumab treatment. **l:** NK2/NK1 ratio of patients with no-response (NR: PD) versus response (R: SD + PR) before and after with ICI. **m:** WT mice were engrafted with 0.5 mio B16-HER2 (s.c.). Mice were treated with AdV5-CCL5 (p.t.) and/or anti-PD-1 (i.p.) antibodies on day 11, 13, 15 and 18 (tumor size 30–70 mm^3^) as indicated. Tumor growth curves are shown. n= 6 mice per condition. *p < 0.05, **p < 0.01, ***p < 0.001, ****p < 0.0001. Error bar values represent SD or SEM (tumor growth curves). Paired 2-tailed Student’s t test was used. For comparisons between three or more groups, one-way ANOVA with multiple comparisons was used. Two-way ANOVA was used to compare tumor growth curves.

To further investigate this possibility, we utilized the patient-derived tumor fragment platform, which allowed us to dissect the early immunological response of human tumor tissue to PD-1 blockade ^32^. We measured IFNγ and CCL5 in the supernatants of tumor fragments from cancer patients exposed to anti-PD-1 (Figure 5e). In line with the correlative analysis of clinical trial data (Figure 5b), we found increased CCL5 in responding fragments (IFNγ upregulation) after anti-PD-1 blockade (Figure 5 f–g).

We next co-cultured human TILs with OVCAR3 cells exposed to anti-PD-1 (Figure 5h). Anti-PD-1 treatment resulted in high concentrations of IFNγ and CCL5 in the supernatants and enhanced killing of OVCAR3 cells (Figure 5i–k). The proportion of CCL5+ cells was higher in the CD49a+ CD16-trNK cells compared to cNK cells in co-cultures exposed to anti-PD-1 (Figure 5k), which confirms trNK cells as the main source of immunotherapy-induced CCL5.

To further underline the beneficial response of CCL5-producing, tissue-resident NK2 cells upon PD-1 blockade, we calculated the ratio of NK2 versus NK1 signature scores for each NK cell in a scRNASeq data set of melanoma patients treated with immune-checkpoint inhibitors (ICI) as a surrogate for NK2 characteristics while controlling for NK cell abundancy. Indeed, we observed that the NK2 phenotype was significantly enriched in patients showing clinical responses (Figure 5l).

Finally, we asked whether CCL5 can further boost anti-PD-1 therapy in trNK cell^poor^ tumors. We combined anti-PD-1 therapy with the adenoviral vector encoding CCL5 in B16-HER2 bearing mice (Figure 5m). While AdV5-CCL5 single treatment did not show any therapeutic efficacy, we observed synergistic therapeutic effects when combined with anti-PD-1 treatment.

Taken together, these data demonstrate that CCL5 production by trNK cells enhances the response to PD-1 blockade, while CCL5-supplementing therapies can enhance its efficacy.

## Discussion

Multimodal immunotherapy combinations that target diverse immune-tumor interactions have become a cornerstone in the therapeutic management of patients with different cancer types and are being extensively explored to maximize the clinical benefit of cancer immunotherapies ^37–40^. Here, we identify the lack of trNK cells as an unrecognized barrier to treatment effectiveness of targeted IL-12 and anti-PD-1 therapy. We show that IL-12 enhanced anti-tumorigenic DC-CD8 T cell interactions which relied on trNK cell-specific induction of CCL5. In tumor models with a limited number of trNK cells and thus low CCL5 levels, only moderate IL-12 responses were observed. However, responses of trNK cellpoor tumors could be rescued by concomitant administration of a CCL5-encoding adenoviral vector which – after converting tumor cells into CCL5 production sites – induced cDC1 infiltration and thus increased DC-CD8 T cell interactions (Supplementary Fig. S8). Due to the unique role of cDC1s in the initiation of T cell responses, both de novo and upon anti-PD-1 checkpoint inhibition, we subsequently observed that CCL5 produced by trNK cells drives the response to anti-PD-1 therapy, which can be delivered using the AdV5 platform. These findings could be utilized to improve immunotherapy by fine-tuning the crosstalk between lymphoid and myeloid immune compartments utilizing multimodal combination immunotherapies adjusted to the pre-existing tumor microenvironment of each patient.

Viral vectors have been shown to be a suitable tool for local immunotherapy, reducing the systemic spread of therapeutic agents and consequently avoiding systemic side effects ^41^. Our adenoviral platform utilizes exogenously added retargeting adaptors consisting of DARPins ^42^. This strategy has unique advantages, compared to targeting by genetic modifications, including the large existing library of DARPins and the rapid selection of new DARPin adapters against any given surface protein ^43,44^. Here, we targeted the model tumor-antigen HER2 which is overexpressed in different cancer types ^45^. To further broaden the clinical applicability, DARPins may be selected to specifically recognize targets on cells other than tumor cells, such as fibroblast activation protein (FAP) on tumor-associated fibroblasts ^46,47^. Recently, the development of high-capacity, helper-dependent AdVs has enabled the expression of transgenes of up to 36 kilobase pairs. This, in combination with the targeted and shielded strategy, has increased the potential of AdVs as an ideal vector for multimodal cancer immunotherapy ^22,48^. Previous work has shown that adenoviral vectors can act as a “self-adjuvants”, allowing the stimulation of multiple innate immune signaling pathways such as toll-like receptors and the induction of type I interferons upon viral entry ^49–51^. While these effects may explain the effect of the empty AdV5-control in tumor delay, the improved anti-tumor immunity supports the potential advantage of adenoviral vectors compared to other gene delivery vectors ^52,53^.

A variety of factors and cell types in the tumor microenvironment underpin the clinical success of cancer immunotherapies, and untangling these complex interactions is critical to understand and improve therapeutic efficacy ^54^. Although typically rare, dendritic cells play a key role in orchestrating anti-tumor immunity. DCs include distinct subsets with non-overlapping functions that can be harnessed for cancer immunotherapy ^33^. Intra-tumoral presence of DCs and particularly the production of IL-12 has been associated with better survival in various cancer types and is positively correlated with clinical outcome to anti-PD-1 therapy ^2^. Consequently, IL-12 has been extensively investigated for use in cancer immunotherapy. Many previous attempts to develop IL-12-based therapies for use in humans, however, resulted in severe toxicities and limited response rates even in tumor-targeted approaches ^14,55^. We here demonstrate that IL-12, provided intra-tumorally by paracrine AdV5 delivery, is capable to bridge innate and adaptive immunity. As a consequence of IL-12 produced by tumor cells after treatment with AdV5-IL12, CD8 T cells, derived from tumors of cancer patients and in tumor models, increased their activation and cytotoxic potential. In the latter, early tumor growth control is achieved by the steady-state of lymphocytes present within the tumor. Yet, to achieve long-term tumor rejection, lymphocyte- and antigen-trafficking to draining lymph nodes is needed. In agreement, we observed increased numbers of NK cells and CD8 T cells in close proximity to blood vessels which suggests enhanced trafficking from the periphery to the tumor. This may be induced by the observed CCL5 secretion by trNK cells or other IFNγ-induced chemokines such as CXCL9 and CXCL10. Moreover, the fact that cDC1s have been described as the main source of CXCL9 and CXCL10 ^56^ could subsequently explain the increased interactions between CD8 T cells and DCs, which have initially been attracted to the TME by CCL5. Consequently, we demonstrated that AdV5-IL12 not only improves lymphocyte homing and activation in the primary tumor but also promotes abscopal anti-tumor effects at distant tumor sides.

NK cells contribute to various immune functions during cancer initiation and progression, including the recognition of cells undergoing stress or early transformation and the direct killing of sensitive cells ^4^. In addition to rapid degranulation upon target recognition, NK cells are powerful producers of a broad variety of pro-inflammatory cytokines and chemokines shaping the inflammatory milieu of tumors ^2,9^. There is growing evidence that the magnitude of NK cell infiltration has a strong prognostic value in various cancers, and predicts clinical outcomes to immune checkpoint blockade in different tumor types ^57^. Investigations of NK cells have originally emphasized their cytotoxic potential, in particular in the context of tumor immunity. Mature potent cytolytic effector CD56^dim^CD16^high^ cNK cells, rapidly secreting pro-inflammatory cytokines and cytotoxic mediators upon receptor-mediated activation, have therefore been considered as the main subpopulations mediating tumor immunity. trNKs in contrast show reduced cytotoxic potential while being specialized on cytokine production such as TNFα, GM-CSF and IL-2 secretion. They have been described to express increased levels of inhibitory checkpoint receptors and were therefore rather associated with tissue homeostasis ^58^. In the context of cancer, they are poorly characterized and controversially discussed. They have been associated with poor survival in human hepatocellular carcinoma, while, on the other hand, they are also reported to join forces with cNK cells to control liver metastasis ^59,60^. We here demonstrate that trNK cells and their expression of CCL5 were essential for the efficacy of AdV5-IL12 and furthermore correlated with response to anti-PD-1 in human melanoma patients. These data uncover a crucial role of trNK cells in priming the tumor immune microenvironment by inducing DC-CD8 T cell interactions and provide direct evidence that the lack of intra-tumoral cDC1 recruitment by trNK cells represents a major barrier for T cell-based therapies. However, further work will need to be performed to comprehensively characterize trNK cells in human tumors, specifically pre- and post-immunotherapy. Moreover, niche-dependent or tumor-derived factors need to be identified that mediate resistance by limiting the accumulation, differentiation, survival, and function of these cells within the tumor microenvironment. Such factors may serve as predictive markers for T cell-focused therapies and define the need for potential combinations with cDC1 attracting agents. With regard to the latter, we here identify CCL5, which we show to improve the efficacy of IFNγ-inducing therapies, such as IL-12 or checkpoint inhibition in tumors with low numbers of trNK cells.

Some controversy exists regarding the role of CCL5 in cancer. A number of studies suggest that CCL5 has potential tumor-promoting effects, either by directly affecting tumor growth ^61^, fostering an immunosuppressive tumor microenvironment ^62^, enhancing tumor cell migration ^63^, or expanding cancer stem cells ^64^. In contrast, other studies show delayed tumor growth and prolonged survival in mouse models with CCL5-expressing tumor cells, as well as a correlation between CCL5 expression and a T cell-inflamed phenotype in cancer patients ^65–67^. In our models, AdV5-CCL5 did not exhibit any tumor-promoting effects but also failed to show therapeutic benefits as a monotherapy. Notably, in agreement with previous work ^9^, AdV5-CCL5 treatment induced cDC1 recruitment. However, only the combination with AdV5-IL12, led to higher co-stimulatory potential and subsequently increased tumor-reactive T cells. This suggests that the anti-tumorigenic properties of CCL5 are likely context-dependent: in the presence of other anti-tumorigenic signals induced by IL-12 or anti-PD1 treatment - which are known to tip the balance to a pro-inflammatory environment - CCL5 may further boost these effects which then translates into improved responses as evidenced by our *in vivo* data and analysis of patient cohorts. Besides CCL5, XCL1 has been described to have similar capacities in recruiting cDC1s to the tumor bed while not being reported to have tumor-promoting capacities ^9^. On this note, XCL1 might be an intuitive alternative payload to guide cDC1s into tumors to improve anti-tumor immunity.

Taken together, our data highlight the importance of trNK cells and their capability to induce T cell immunity by enhancing DC-T cell crosstalk in IFNγ inducing therapies and may inform novel combination strategies utilizing viral vector platforms as an approach to further potentiate this NK cell-DC-T cell crosstalk. Our data highlight a relevant tumor-eliminating positive feedback mechanism to be prioritized for clinical development, particularly in patients with immune-excluded and/or resistant tumors.

## Methods

### Redesign of AdV5 shuttle vector

The pShuttle vector from the AdEasy™ Adenoviral Vector System (Agilent Technologies) was redesigned to allow for the rapid generation and exchange of modular expression cassettes encoding a variety of payloads ^68^. The multiple cloning site (MCS) of the pShuttle vector was replaced with synthetic MCS modules, called MCS1 or MCS2, via Gibson Assembly (New England Biolabs). The synthetic MCS1 module contained, from 5’ to 3’, the CMV promoter, NheI restriction site, XhoI restriction site, and the polyA site from BGH as previously described 20. The MCS2 module contained, from 5’ to 3’, the SV40 promoter, SpeI restriction site, SalI restriction site and the polyA site from SV40. MCS modules were synthesized by GeneArt (Life Technologies Europe BV) containing the N-terminal flanking DNA 5’-GAA TAA GAG GAA GTG AAA TCT GAA TAA TTT TGT GTT ACT CAT AGC GCG TAA -3,’ and C-terminal flanking DNA 5’-TAA GGG TGG GAA AGA ATA TAT AAG GTG GGG GTC -3’ for Gibson Assembly into pShuttle to generate the plasmids pShuttle-MCS1 or pShuttle-MCS2. Additionally, a plasmid called pShuttle-MCS1-MCS2 was constructed where both the MCS1 and MCS2 modules were inserted in tandem into the same pShuttle construct.

### Construction of payload construct

Murine and human IL-12 constructs were generated from translated Genbank cDNA sequences for the IL-12B/p40 (NCBI Reference: BC103608.1 or BC074723.2, respectively) and IL-12A/p35 (NCBI Reference: BC146595.1 or BC104984.1, respectively) connected by a F2A peptide as previously described ^69^. The murine CCL5 gene was generated from Uniprot sequence P30882. Cytokines included their native signal sequences, were codon-optimized for human or mouse expression, respectively, and synthesized by GeneArt (Thermo Fisher Scientific). For reporter assays, a firefly luciferase reporter was synthesized from GenBank: BAL46512.1. Payload constructs were inserted into redesigned pShuttle vectors by Gibson assembly or standard ligation cloning to generate pShuttle-MCS1 including the payload. The pShuttle-MCS1 backbone (without payload) was used to generate the AdV5-control vector. In general, pShuttle-MCS1-IL12 (AdV5-IL12), pShuttle-MCS1-CCL5 (AdV5-CCL5), and pShuttle-MCS1-Luciferase (AdV5-Luc) were used to generate immunotherapeutic vectors. For the combinatorial approach optimization (Figure S5), pShuttle-MCS2-CCL5 (AdV5-SV40-CCL5) and pShuttle-MCS2-IL12 (AdV5-SV40-IL12), pShuttle-MCS1-IL12-MCS2-CCL5 (AdV5-CMV-IL12-SV40-CCL5) and pShuttle-MCS1-CCL5-MCS2-IL12 (AdV5-CMV-CCL5-SV40-IL12) were used.

### Virus production

The plasmid containing the adenoviral genome, pAdEasy-1, from the AdEasy™ Adenoviral Vector System (Agilent Technologies) was previously modified to include a mutation to the hypervariable loop 7 (HVR7) of the hexon, which prevents blood factor X binding to virions and thus reduces liver infection ^22^. To generate viral constructs, the modified pAdEasy-1_HVR7 plasmid was co-transformed with the pShuttle-MCS variants listed above into recA-proficient E. coli BJ5183 cells, from which the desired recombinants, obtained by homologous recombination, could be isolated for virus production. Packaging and amplification of adenoviral particles was performed by Vector Biolabs (Malvern, PA, USA) and they were purified on two consecutive cesium chloride density gradients and provided directly in PBS with 5% glycerol.

### Protein purification of adenoviral shield and retargeting adaptor

The human HER2 adenoviral retargeting adapter (G3_1D3nc_SHP1) was expressed and purified as previously described ^22,42^. Endotoxin was removed from purified adapters using the Endotrap^®^ HD Endotoxin Removal System (Hyglos GmbH) and adapters were stored at −80°C in endotoxin-free Dulbecco’s PBS (Millipore TMS-012-A). The adenoviral shield was purified in Sf9 insect cells as previously described ^22^.

### Mice

C57BL/6 and Balb/c mice were bred in-house at University Hospital Basel, Switzerland. Batf3 KO (B6.129S(C)-Batf3<tm1Kmm>/J) mice were obtained from the Jackson laboratory, USA. Animals were housed under specific pathogen-free conditions. All animal experiments were performed in accordance with Swiss federal regulations. Sex-matched littermates at 8-12weeks of age at the start of experiments were used.

### Tumor models

C57BL/6 and Batf3KO mice were injected subcutaneously into the right flank with 0.5 mio syngeneic murine B16 D5 melanoma cells expressing HER2 (kindly provided by Dr. L. Weiner, Georgetown University, Washington, DC) suspended in phenol red-free DMEM (without additives). EMT6 murine breast cancer cells expressing HER2 (1 mio) were injected into the mammary gland of female Balb/c mice ^70^. Cell lines were tested for mycoplasma contamination before injection. Tumor volume was calculated according to the formula: D/2 x d x d, with D and d being the longest and shortest tumor diameter in mm, respectively.

### Immunotherapy treatments

Tumor bearing mice, with a tumor size of approximately 30-70 mm^3^, were treated with each 1.5×10^8^ PFU of HER2-targeted and shielded adenoviral vectors in 50 μl of PBS (peritumorally), and/or 12.5 mg/kg mouse anti-PD-1 (RPM1–14, BioXCell) or left untreated. For depletion studies, CD8 T cells were depleted by administering anti-CD8a (53-6.72, BioXCell) at 10mg/kg (i.p.) once per week. NK depletion was performed by administering anti-Asialo-GM1 (Poly21460, Biolegend) 50 μl (i.p.) in Balb/c mice or anti-NK1.1 (PK136, BioXCell) 10 mg/kg (i.p.) in C57BL/6 mice every 4-5 days. IFNγ neutralization was performed using anti-IFNγ antibody (XMG1.2, BioXcell) at 25 mg/kg in 200 μl PBS injected every 2-3 days. To neutralize CCL5, we injected 32 μg anti-CCL5 antibody (500-P118, Peprotech) in 200 μl PBS per mouse intraperitoneally. Depletion and neutralization schedules were started the day before immunotherapy treatment unless stated otherwise.

### Bilateral tumor models

Balb/c mice were inoculated with 1 mio EMT6-HER2 tumor cells (i.m.) in the right flank. Four days later, 0.25 mio EMT6 wt cells were injected into the contralateral site (i.m.). On days 7, 9, 11, and 13 post first tumor inoculation, EMT6-HER2 tumors were treated with each 1.5×10^8^ PFU of HER2-targeted and shielded adenoviral vector encoding IL-12 in 50 μl of PBS (peritumorally). Tumor volumes of contralateral (EMT6 wt) tumors were measured.

### Tumor re-challenge

Long-term surviving mice from AdV5-IL12 therapy were re-challenged with EMT6 wt and EMT6-HER2 tumors in each flank 60 days after primary tumor rejection. EMT6 wt and EMT6-HER2 re-challenge doses were 0.25 mio cells and 1 mio cells, respectively. As a control, naive Balb/c mice were implanted alongside re-challenged mice.

### FTY720 treatments

Mice were implanted with EMT6-HER2 or B16-HER2 tumors intramammary or subcutaneously, respectively. Mice were treated or not with 1.25 mg/kg of FTY720 (Cayman Chemical) i.p. daily throughout the duration of the experiment. Injections were started one day before tumor inoculation or the day before adenoviral treatment.

### *In vivo* pharmacokinetic experiments

Mice were implanted with EMT6-HER2 or B16-HER2 tumors intramammarily or subcutaneously, respectively. Once the tumors reached an average volume of 30-70 mm^3^, luciferase-encoding retargeted and shielded AdV5 (1.5×10^8^ PFU per mouse; AdV5-Luc) were injected peritumorally. The luciferase signal was determined in live animals one day after virus injection and 10 minutes after intraperitoneal injection of 150 mg/kg D-luciferin (PerkinElmer) using the *in vivo* imaging system NightOWL II LB 983 (Berthold) over two weeks. Following live imaging, luciferase activity was determined in isolated tumors and organs (draining and non-draining lymph nodes, spleen, liver, kidney, lung, and heart). The overlay of the real image and the luminescence representation allowed the localization and measurement of luminescence emitted from xenografts. The signal intensities from manually derived regions of interest (ROI) were obtained and data were expressed as photon flux (photon/s). All measurements were performed under the same conditions, including camera settings, exposure time (60 s), distance from lenses to the animals and ROI size.

### Multiparameter flow cytometry

Tumor tissue was isolated from mice, weighed, and minced using razor blades. Tissue was then digested using accutase (PAA), collagenase IV (Worthington), hyaluronidase (Sigma), and DNAse type IV (Sigma) for 60 min at 37°C with constant shaking. The cell suspensions were filtered using a cell strainer (70 μm). Precision Counting beads (Biolegend) were added before staining to quantify the number of cells per gram tumor. Single cell suspensions were blocked with rat anti-mouse FcγIII/II receptor (CD16/CD32) blocking antibodies (“Fc-Block”) and stained with live/dead cell-exclusion dye (Zombie UV dye; Biolegend). The cells were then incubated with fluorophore-conjugated antibodies directed against cell surface antigens, washed, and resuspended in FACS buffer (PBS + 2% FBS). For intracellular/intranuclear antigens, cells stained with cell surface antibodies were fixed and permeabilized using Foxp3/transcription factor staining buffer set (eBioscience) prior to incubation with antibodies directed against intracellular antigens. Cell populations were analyzed on a Cytek Aurora and Cytoflex.

### Bioinformatic analysis of flow cytometry data

FCS files containing pre-gated alive CD45+ single cells were read into R using the flowCore package (flowCore: flowCore: Basic structures for flow cytometry data version 2.2.0 from Bioconductor). A logicle transform was performed per channel, with parameters calculated from aggregated data from all samples. CD45 low cells with a transformed value of < 2.5 were removed from further analysis (threshold set on the left side of trough of density plot of transformed CD45 values from all samples). The measurements were randomly subsampled to 1.5 x 10^5^ cells per condition to expedite downstream operations. Principal Component Analysis (PCA) was performed using all markers except for CD45, Live-Dead, FSC*, and SSC*. Uniform Manifold Approximation and Projection (UMAP) was performed for visualization using the CATALYST module’s runDR function (CATALYST: Cytometry dATa anALYSis Tools version 1.14.0 from Bioconductor). Clustering was performed using Rphenograph (0.99.1), an R implementation of PhenoGraph (GitHub - JinmiaoChenLab/Rphenograph: Rphenograph: R implementation of the PhenoGraph algorithm) ^74^. Five clusters with universally high expression across all markers were removed, and steps starting with PCA were repeated. Three resulting clusters were again removed and the process iterated once more to yield the final clustering, UMAP, and heatmaps ^75^. Supplemental UMAP visualization of scaled marker expression done with CATALYST’s plotDR function. Main cell types were assigned by marker expression. To confirm certain assignments, cell populations were gated using FlowJo (10.6.2) and compared to assigned populations (e.g. NK cells: NKp46+, CD3-, CD19-Ly-6G-, F4/80-).

### Multiparameter fluorescence microscopy

Tumors embedded in OCT were sectioned into 7 μm-thick slices and attached to poly-L-lysine coated square coverslips. Sections were analyzed using CODEX^®^ (Akoya Biosciences), a highly multiplexed imaging platform, which allows the staining of solid tissue sections with a panel of up to 40 antibodies at once ^23^. In brief, CODEX uses a unique DNA barcode system to label each antibody clone individually. These barcodes can be detected by reversible hybridization to their corresponding reporter. The respective reporters, which are conjugated with the fluorophores AF488, Atto550, or Cy5, are applied onto the tissue sections, imaged, and removed in a multicycle experiment. For this purpose, the manufacturer’s protocol "CODEX User Manual Rev A.0" (provided by AKOYA Biosciences) was followed.

The antibody panel was composed of commercially available AKOYA-conjugated antibodies and self-conjugated custom antibodies (CODEX Conjugation Kit, AKOYA Biosciences). Tissue staining was performed with the CODEX Staining Kit (AKOYA Biosciences). Briefly, the tissue was thawed with drierite beads, fixed with acetone, rehydrated, and fixed with 1.6% paraformaldehyde (PFA). After blocking, the tissue was stained with the established antibody panel consisting of 33 barcoded antibodies at the same time. The bound antibodies were fixed to the tissue with 1.6% PFA, ice-cold methanol, and a fixative solution (AKOYA Biosciences).

The inverse microscope DMi8 (Leica) was used for acquisition (20x magnification, xyz acquisition mode 14 Z-stacks each 14.99 μm, "Best Focus" autofocus with default settings). The generated fluorescence data were formatted with the Akoya CodexDriver V2 and subsequently processed using CODEX Processor (Version v1.5.0.48b) or the Kheops PlugIn in ImageJ and QuPath0.2.3 and StarDist ^76–78^. The processing steps included 1) XY Processing with tile registration and shading correction; 2) Z-Stack Processing with deconvolution, drift compensation, overlap cropping, background subtraction (min-min mode), and best focus detection; 3) Stitching with best focus interpolation and tile overlap of 10% and 4) Cell segmentation on the nuclear stain with the radius 8 or threshold (0.69), channels (’DAPI’), normalize percentiles (1.99), pixel size (0.423), cell Expansion (2.8), cell constrain scale (1.5) in StarDist, respectively.

The processed and segmented data were analyzed with the CODEX Multiple Analysis Viewer (Version 1.2.0.297) or the PhenoGraph algorithm using R 4.0.2. Manual gating was performed to distinguish between CD45+ cells (immune cells) and CD45-cells (tumor and stroma cells). Clustering was performed with VORTEX using unbiased hierarchical X-shift clustering (K = 55 resulting in 81 immune clusters) ^79^. Clusters were manually verified and assigned to main immune cell populations using CODEX Multiple Analysis Viewer or QuPath. Subsequently, mean marker expression and interaction counts between cellular main populations (cells with <50 μm proximity) were determined.

### Interaction analysis

Interaction counts between main cell populations (contact defined as <50 μm proximity) were used for further analysis. The cluster “L” containing lymph vessels was excluded from further analysis, because of the very distinct localization of its cells as small vessels. Expected interaction counts for each cell-cell interaction pair were calculated as:

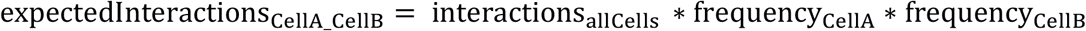

Next, a negative-binomial generalized linear model for the interaction counts with an offset of log(expectedInteractions) was generated in R 4.0.2. The mathematical interaction terms between cell-cell type comparison (e.g. DC_vs_CD8) and experimental condition (untreated, AdV5-empty, AdV5-IL12) were used to calculate odds ratios and p-values for changes in cell-cell type interactions between experimental conditions. Odds ratios for different subsets of immune cells were plotted using ggplot2.

### Intra-tumoral and systemic cytokine measurements

Serum was collected in EDTA containing tubes (Sarstedt) and IL-12 levels were determined using the IL-12 p70 Mouse Uncoated ELISA Kit (Invitrogen). Isolated tumors were snap-frozen on dry ice. Before thawing, a 5 mm metal bead and 1 ml of lysis buffer (20 mM Tris HCl (pH 7.5), 0.5% Tween 20, 150 mM NaCl, Sigma protease inhibitors 1:100) were added to the tubes. Tumors were lysed using a TissueLyser (Qiagen) for 5 min at 25 Hz. After centrifugation, protein concentrations were determined with the Pierce BCA Protein Assay Kit (Thermo Scientific). CCL5 concentration in the tumor lysates was analyzed by ELISA (Mouse RANTES Uncoated ELISA Kit Invitrogen) and normalized to the determined total protein concentrations.

### Patients and sample preparation

Surgical specimens were mechanically dissociated, digested with accutase (PAA Laboratories), collagenase IV (Worthington), hyaluronidase (MilliporeSigma), and DNAse type I (MilliporeSigma), filtered, washed, and frozen as single cell suspension for future use. For human *ex vivo* tumor cultures, surgical specimens were dissected into tumor fragments and frozen for future use. Human PBMCs were isolated by density gradient centrifugation using Histopaque-1077 (MilliporeSigma) from buffy coats obtained from healthy blood donors (Blood Bank, University Hospital Basel). PBMCs were frozen for later use in liquid nitrogen. Ethics approval was obtained from the local ethical committee to analyze the tissue and blood samples (Ethikkommission Nordwestschweiz) and written informed consent was obtained from all patients prior to sample collection.

### *Ex vivo* human immune cells and tumor co-culture experiments

Healthy donor PBMCs or tumor digest samples, processed as described above, were co-cultured with OVCAR3 human ovarian cancer cell line, similarly to as described in Natoli et al. 2020. Briefly, 6000 OVCAR3 cells were seeded on the wells of a flat-bottom 96-well plate in RPMI containing L-glutamine (R8758, Sigma-Aldrich), supplemented with penicillin/streptomycin (100 ng/ml, Sigma-Aldrich) and 10% FBS (Sigma-Aldrich). After 2 hours, the medium was replaced with fully supplemented RPMI containing AdV5-hu-IL12 or empty vector control (AdV5-control) at a PFU of 1000/cell. Additional control wells were left untreated. After 2 hours of incubation, the medium was replaced to remove the virus and the tumor cells were incubated for 2 days at 37°C, 5% CO2. 300,000 healthy donor PBMCs or single cells from tumor digest samples were then added to the wells of the 96-well plate in a final volume of 200 μl per well. Tumor cell (OVCAR3) viability was assessed by an MTT assay and flow cytometry was conducted on the suspension cells (PBMCs, TILs) after 3 days of co-culture. The supernatant was collected after 6 days of co-culture to assess IFNγ and CCL5 levels using a Human IFNγ ELISA Set (BD OptEIA, 555142) and ELISA MAX™ Deluxe Set Human CCL5 (Biolegend, 440804), respectively, according to the manufacturers’ instructions.

### *Ex vivo* tumor fragment culture

Tumor fragment cultures were prepared as described in Voabli et al. 2021. Briefly, frozen patient tumor fragments were slowly thawed at 37°C and extensively washed in PBS and warm RPMI medium containing L-glutamine (R8758, Sigma-Aldrich), supplemented with penicillin/streptomycin (100 ng/ml, Sigma-Aldrich), 10% FBS (Sigma-Aldrich), 1x MEM Non-Essential Amino Acids (Gibco) and 1 mM sodium pyruvate (Sigma-Aldrich).

Single tumor fragments were then embedded in a total of 80 μl of an artificial extracellular matrix within the wells of a flat-bottom 96-well plate. The extracellular matrix was prepared by mixing ice-cold sodium bicarbonate (Sigma; 1.1% final concentration), collagen I (Corning; 1 mg/ml final concentration) and fully supplemented RPMI with ice-cold matrigel (Matrix High Concentration, Phenol Red-Free, BD Biosciences; 4 mg/ml final concentration). 40 μl of matrix was solidified by incubation at 37°C for 20-30 min. One tumor fragment per well was placed on top of the pre-solidified matrix, after which a second layer of 40 μl matrix was added. Plates were then placed in a 37°C incubator for further 20-30 min. 110 μl of fully supplemented RPMI was added on top of the matrix. 1 mio PFU of AdV5-hu-IL12 or control AdV5-control or nivolumab (10 μg/ml final concentration) were added to each well containing individual tumor fragments. Between 6-12 fragments were used for each treatment conditions. After 48 hours of incubation at 37°C, the fragments were pooled and enzymatically digested and filtered into single cell suspensions, as described above. Flow cytometry was conducted and the supernatant was collected from each well to assess IFNγ and CCL5 levels using a Human IFNγ ELISA Set (BD OptEIA, 555142) and ELISA MAX™ Deluxe Set Human CCL5 (Biolegend, 440804), respectively, according to the manufacturers’ instructions.

### MTT assay

To assess tumor cell viability in co-culture experiments, an MTT assay was used as follows. The medium from the co-culture wells was removed and the wells were gently washed once with PBS to remove suspension cells. MTT (Sigma-Aldrich) was then added at 500 μg/ml and the tumor cells were incubated at 37°C for 2–3 hours. Formazan crystals were resuspended in 90 μl of dimethyl sulfoxide (DMSO; Sigma-Aldrich) and absorbance was measured at a wavelength of 570 nm.

### Bioinformatic analysis of published gene expression data of human melanoma samples

We received the normalized Nanostring RNA expression data from the NCT01502293 trial (Oncosec), in which patients received IL-12-encoding mRNA by intra-tumoral electroporation ^16^. The gene signatures for different immune sub-populations were retrieved from the PanCancerImmunology Nanostring panel. The transcripts within the “Cytotoxic cells” signature can’t be attributed to a single cell type and are found in other signature lists of both NK and CD8 T cells. Therefore, its transcripts were merged with the cytotoxic NK and CD8 T cells resulting in the “CD8+ T cells” and “NK cells” signatures. Importantly, neither “CD8+ T cells” and “NK cells” contained the *CCL5* transcript. Downstream analysis of all Nanostring data was performed using R version 4.0.2 and visualized in GraphPad Prism 9.

Immune cell infiltration was estimated by calculating a signature score as described by Cursons et al. 2019. Briefly, all transcripts for each sample were ordered by decreasing expression and the signature score was defined as:

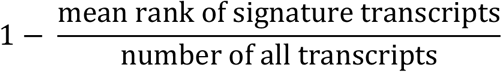

Thus, a high signature score indicates enrichment of signature transcripts among genes with high expression. To find genes that correlate with the NK cells signature score, we performed a linear regression for signature scores with log-transformed transcript counts. p-values were adjusted by the Benjamini-Hochberg correction method.

We also reanalyzed RNAseq data of samples from tumor patients treated with anti-PD-1 antibodies (nivolumab) ^36^. Data was downloaded from the GEO under accession number GSE91061. Only patients with both pre- and post-treatment samples and evaluated responses were analyzed. Counts were normalized by library size using edgeR and displayed as log counts per million using GraphPad Prism 9. Paired two-way ANOVA with multiple comparisons and the post-hoc test were used to calculate significances for increased *CCL5* expression upon treatment in CB. The availability of RNAseq data allowed the use of more complex gene signatures than with Nanostring data above. Therefore, we used the signatures described by Tirosh et al and Cursons et al. for this analysis ^80,81^. As we wanted to correlate signatures with *CCL5* expression, *CCL5* was excluded from the T cell signature provided by Tirosh et al. 2016. Signature scores were calculated as described above.

For the reanalysis of the Zillionis et al. scRNAseq dataset we downloaded the pre-cleaned count matrix from GEO under accession number GSE127465. The dataset was analyzed using the Bioconductor packages SingleCellExperiment and scater. PCA was performed using the top 4000 variable gene log counts. UMAP dimension reduction was performed on the PCA values using the scater runUMAP with 15 neighbors. *CCL5* expression per cell was visualized by ggplot2 using UMAP coordinates. To construct the cNK (NK1) and trNK (NK2) signatures we extracted the z-scores for genes enriched in either cluster using the online tool of the original publication. We then selected genes with a z-score of >0.5 and a minimum difference in z-score between the two subsets of 0.5.

We also obtained signatures for tissue resident NK cells from Marquart et al and a signature for CD56^bright^ CD16^neg^ NK cells from Hanna et al ^34,35^. A score for each signature as above was then calculated for each cell in the Zillionis dataset. Violin plots of these values were then plotted using Graphpad Prism.

Similarly, we also obtained the scRNAseq dataset described in Sade-Feldmann et al. under GEO accession number GSE120575. As described in their original publication, genes were filtered for protein coding genes and min expression of >4.5 logcounts in at least 10 cells. Cells were filtered to have >2.5 mean logcounts of their listed housekeeping genes. Cells with a high fraction mitochondrial reads (>3 standard deviations; dying cells) and very many detected genes (> 4 standard deviations; doublets) were excluded. PCA and UMAP dimension reductions were calculated as above. Clustering was performed using buildSNNGraphs of the scran package from PCA values and k = 15. Cluster8 consisted of NK cells based on its high expression of *NCR1*, *NCAM*, and *FCGR3A*. For each cell in this NK cluster, we calculated the mean score for cNK (NK1) and trNK (NK2) signatures as described above. Then we summarized the mean cNK and trNK value for the NK cells of each patient. The values for responding and non-responding patients were then visualized as a ratio between trNK and cNK cells using Graphpad Prism.

### Statistical Analysis

For normally-distributed datasets, we used 2-tailed Student’s t test and one-way ANOVA followed by Holm-Sidak multiple comparison test. When variables were not normally distributed, we performed non-parametric Mann-Whitney or Kruskal-Wallis tests. For survival analysis, p values were computed using the Log Rank test. Two-way ANOVA was used to compare tumor growth curves and grouped data sets. p values > 0.05 were considered not significant, p values < 0.05 were considered significant. * p value < 0.05, ** p value < 0.01, *** p value < 0.001, **** p value < 0.0001.

## Data availability

The previously published RNAseq and scRNAseq data sets we re-analyzed are accessible at GEO under accession number GSE91061, GSE120575 and GSE127465, respectively. Source data and all other supporting data of this study are available from the corresponding authors on a reasonable request.

## Code availability

Codes used for the analysis of flow cytometry data and multidimensional IF microscopy in Fig. 1, as well as the analysis of published expression data can be obtained from the corresponding authors upon request.

## Acknowledgments

We thank members of the Cancer Immunology and Cancer Immunotherapy Laboratory at the Department of Biomedicine for helpful discussions and suggestions. We are grateful to A. Ignatenco and B. Simic for cloning viral constructs and to F. Weiss, and P. Freitag for producing and providing the biological reagents. We thank Priska Auf der Maur (University Hospital of Basel) for designing the graphical abstract. This work benefited from clinical data provided by OncoSec Medical Incoporated. This work was funded by the Schweizerische Nationalfonds Grant CRSII5_170929 to A.Z. and A.P., by the Schweizerische Nationalfonds Grant 320030_188576/1 to A.Z., by the National Cancer Institute of the National Institutes of Health under award number F32CA189372 (to S.N.S.), by the University of Zurich Forschungskredit 2017 ID 3761 (to D.B.), and by the Cancer League beider Basel Grant KLbB-5325-03-2021 (to N.K.).

## Authors contributions

N.K. designed the study, performed experiments, analyzed data, and wrote the manuscript. M.P.T., M.N., F.W., V.K., M.B., and D.S.T. planned and performed experiments and generated and analyzed data. D.B., S.N.S., D.B., K.P.H, and P.Z. designed, cloned, or produced biological reagents. D.S. and E.B. processed and analyzed data. M.P.T., M.N., D.B., S.N.S., H.L., J.B., M.A.S., A.S.K., and A.P. provided input for research design and interpretation and edited the manuscript. A.Z. directed the study and wrote the manuscript. All authors reviewed and approved the manuscript.

## Notes

**Conflict of interest statement** The authors declare no potential conflicts of interest.

### Competing Interest Statement

The authors have declared no competing interest.

