## Supplementary Figures for "Adaptive anti-tumor immunity is orchestrated by a population of CCL5-producing tissue-resident NK cells"

### Supplementary Information

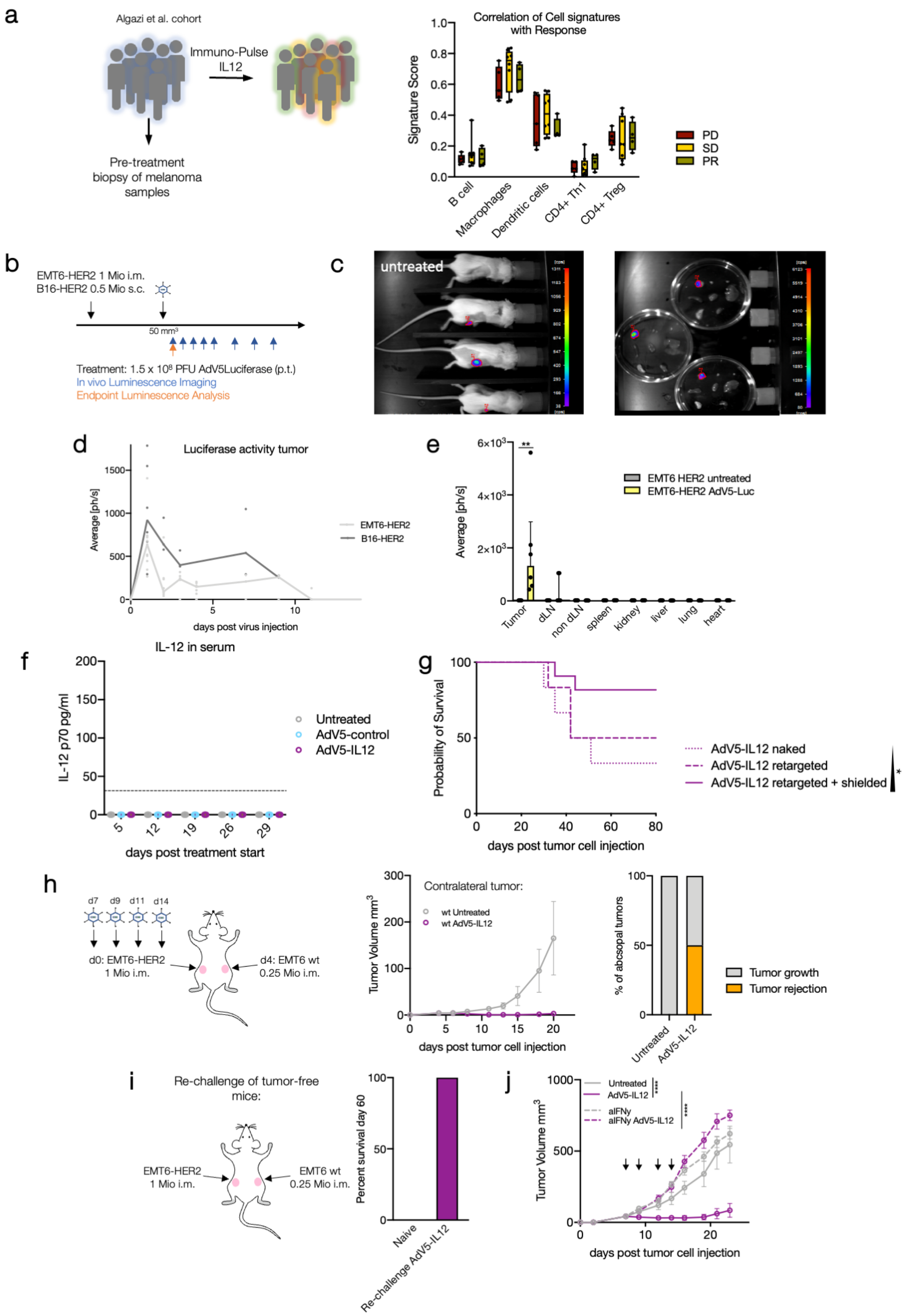


**Supplementary Fig. S1: Related to Figure 1**

**a:** Cell signature scores measured by Nanostring in skin tumor biopsies from 19 melanoma patients before intra-tumoral treatment with ImmunoPulse IL-12 were correlated with clinical response (PD: progressive disease, SD: stable disease, PR: partial response) **b-e:** Wildtype (WT) mice were engrafted with 1 mio EMT6-HER2 (i.m.) or 0.5 mio B16-HER2 (s.c.). Mice were p.t. treated with AdV5-Luciferase on day 7 or day 11 (tumor size 30–70 $mm^3$), respectively. The luciferase signal was live imaged at the indicated time points (blue arrows; day 1, 2, 3, 4, 5, 7, 9, 11 and 13 post virus injection). Representative luciferase signal in three treated and one untreated animal one day post AdV5-Luciferase injection are shown. After isolation of tumor, draining lymph node, non-draining lymph node, spleen, kidney, liver, heart and lung, luciferase signal was measured again. **d:** Quantification of in vivo luciferase signal in EMT6-HER2 (light grey) and B16-HER2 (dark grey) tumors. **e:** Quantification of luciferase signal one day post AdV5-Luciferase treatment in isolated indicated organs of EMT6-HER2 bearing mice. **b-e:** n= 6 mice per condition. **f:** Wildtype (WT) mice were engrafted with 1 mio EMT6-HER2 intramammarily (i.m.). From day 7 (tumor size 30–70 $mm^3$), mice were treated with $1.5x10^8$ PFU of HER2-targeted and shielded adenoviral vectors (peritumorally) encoding for IL-12 or empty control cassette (AdV5-control) on day 7, 9, 11 and 14. On the indicated days, serum was collected and IL-12 concentration was determined by ELISA. Dotted line denotes detection limit of the ELISA. **g:** Wildtype (WT) mice were engrafted with 1 mio EMT6-HER2 intramammarily (i.m.). From day 7 (tumor size 30–70 $mm^3$), mice were treated with $1.5x10^8$ PFU of naked, HER2-targeted or HER2-targeted and shielded adenoviral vectors (peritumorally, p.t.) encoding for IL-12 or an empty control cassette (AdV5-control) on days 7, 9, 11 and 14 p.t. Kaplan-Meier survival curves are shown. Black arrows denote days of treatment. n = 6. Log rank test for trend was performed to determine significant changes. **h:** Wildtype (WT) mice were engrafted with 1 mio EMT6-HER2 (i.m.) and with 4 days delay with 0.25 mio EMT6 wt cells on the contralateral flank. EMT6-HER2 tumors were peritumorally treated with AdV5-IL12. Tumor growth of contralateral tumor (EMT6 wt) was measured. Tumor growth curve and percentage of rejected contralateral tumors are shown. n = 6 mice per group. **i:** Mice which rejected tumors after AdV5-IL12 treatment were rechallenged (60d after tumor rejection) with 1 mio EMT6-HER2 i.m. and 0.25 mio EMT6 wt cells (no HER2 transgene) on each flank. Percentage of survival 60d post rechallenge is shown. n = 12, naive mice served as a control. *$p < 0.05$, **$p < 0.01$, ***$p < 0.001$, ****$p < 0.0001$. Error bar values represent SD or SEM (tumor growth curves). For comparisons between three or more groups, one-way ANOVA with multiple comparisons was used. For survival analysis, p values were computed using the Log Rank test. Two-way ANOVA was used to compare tumor growth curves.

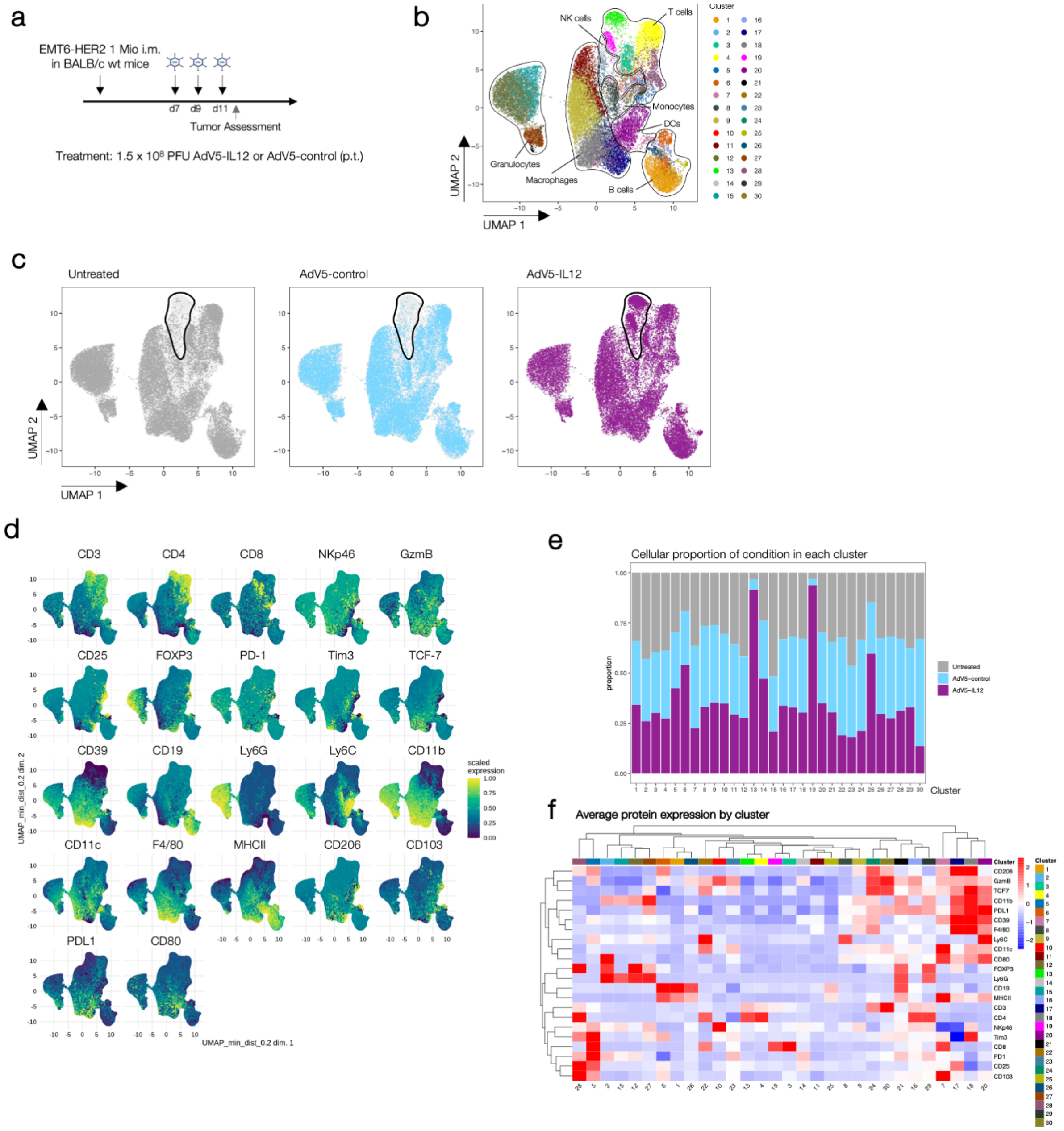

**Supplementary Fig. S2: Related to Figure 1**

**a:** Wildtype (WT) mice were engrafted with 1 mio EMT6-HER2 intramammarily (i.m.). Starting from day 7 (tumor size 30–70 mm$^3$), mice were treated with 1.5x10$^8$ PFU of HER2-targeted and shielded adenoviral vectors (p.t.) encoding for IL-12 or empty control cassette (AdV5-control) on day 7, 9 and 11. On day 12 post inoculation, tumors were isolated and single cell suspensions were analyzed by flow cytometry. **b:** UMAP projection is depicting the alive CD45+ tumor infiltrating lymphocytes colored by cluster. **c:** UMAP projection is showing distribution of cells colored by treatment condition (dark grey: untreated; blue: AdV5-control; magenta: AdV5-IL12). **d:** UMAP-projection of pooled conditions showing expression analyzed proteins supporting cell-type assignments. **e:** Percentage of cells in each cluster by treatment. **c:** Heatmap showing average protein expression by cluster. **f:** Heatmap showing protein expression of 2 x 10$^4$ random cells assigned to conditions (Untreated, AdV5-control or AdV5-IL12) and cluster. n = 5-6 mice per group.

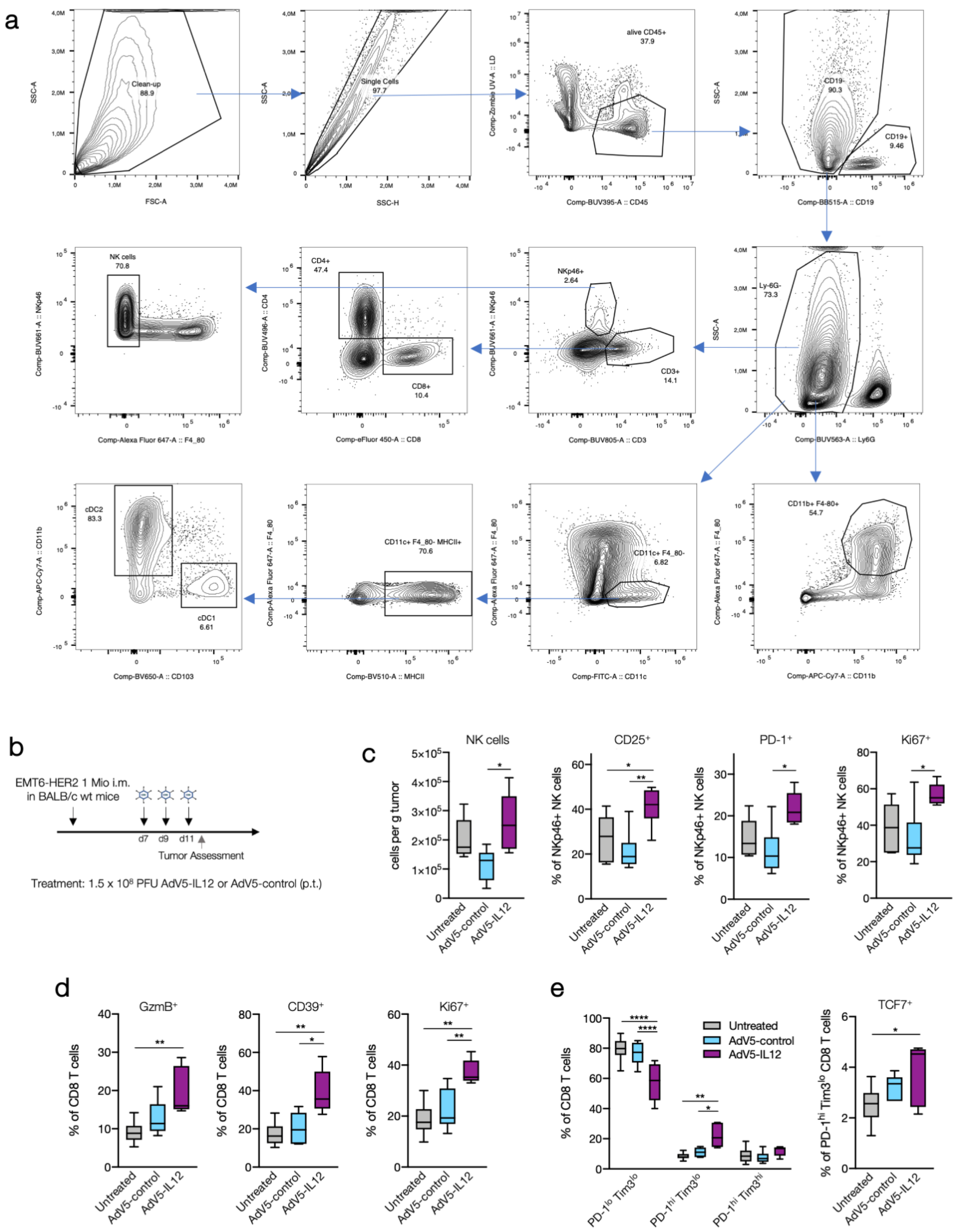

**Supplementary Fig. S3: Related to Figure 1**

**a:** To gate on singlets, SSC-A and SSC-H were used. After gating on alive CD45+ cells, CD19+ B cells and Ly-6G+ Granulocytes were excluded. To define T cells (CD3+), NK cells were excluded (NKp46+ F4/80-) and further distinguished between CD8 T cells and CD4 T cells. Macrophages were defined using F4/80+ and CD11b+. DCs were defined as CD11c+ F4/80- cells. MHCII was used to gate on cDCs. Subsequently to distinguish cDC1s from cDC2s, CD11b and CD103 was used. **b-e:** Wildtype (WT) mice were engrafted with 1 mio EMT6-HER2 intramammarily (i.m.). Starting from day 7 (tumor size 30–70 mm$^3$), mice were treated with 1.5x$10^8$ PFU of HER2-targeted and shielded adenoviral vectors (p.t.) encoding for IL-12 or empty control cassette (AdV5-control) on day 7, 9 and 11. On day 12 post inoculation, tumors were isolated and single cell suspensions were analyzed by flow cytometry. **c:** Quantification of NK cells (NKp46+, CD3-, Ly6G-, CD19-, F4/80-) per gram tumor and proportion of CD25+, PD-1+ or Ki67+ of NK cells between the different treatment conditions. **d-e:** Proportion of granzyme B+ (GzmB+), CD39+ or Ki67+ of CD8 T cells (CD3+, CD4-, NKp46-, CD19-) between the different treatment conditions. Proportion of PD-$1^{lo}$TIM$3^{lo}$, PD-$1^{hi}$TIM$3^{lo}$ or PD-$1^{hi}$TIM$3^{hi}$ intra-tumoral CD8 T cells in each treatment group. n = 5-6 mice per group. *$p < 0.05$, **$p < 0.01$, ***$p < 0.001$, ****$p < 0.0001$. Error bar values represent SD. For comparisons between three or more groups, one-way ANOVA with multiple comparisons was used.

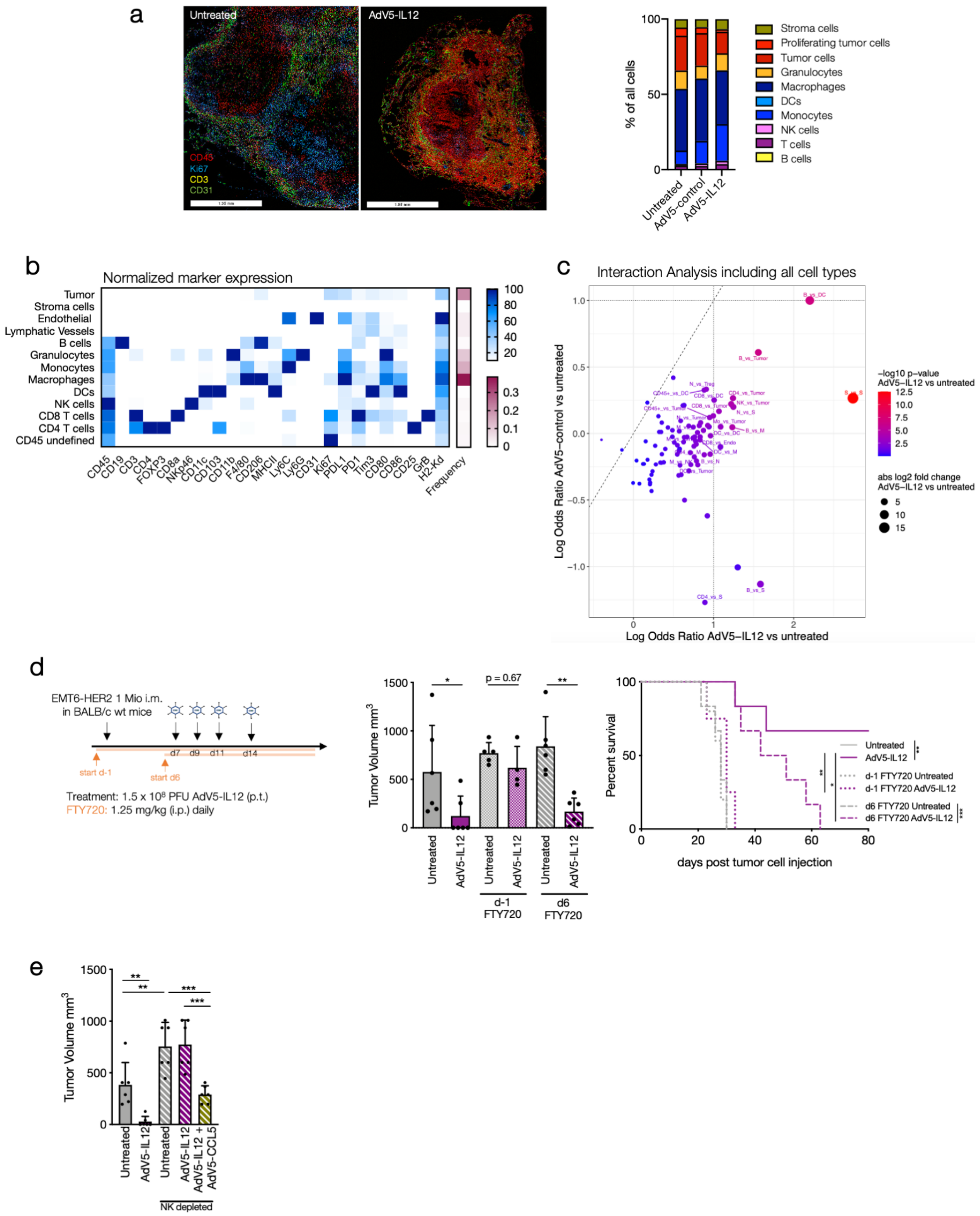


**Supplementary Fig. S4: Related to Figure 1 and 2**

**a-b:** Wildtype (WT) mice were engrafted with 1 mio EMT6-HER2 intramammarily (i.m.). Starting from day 7 (tumor size 30–70 mm$^3$), mice were treated with 1.5x10$^8$ PFU of HER2-targeted and shielded adenoviral vectors (peritumorally) encoding for IL-12 or empty control cassette (AdV5-control) on day 7, 9 and 11. On day 12 post inoculation, tumors were isolated, embedded in OCT and analyzed by multiparameter immuno-fluorescence microscopy. n= 3 mice per condition. **a:** Representative IF pictures are showing untreated and AdV5-IL12 treated tumors (CD45: red, Ki67: blue, CD3: yellow, CD31: green) including quantification of main clusters between conditions. **b:** Heatmap showing normalized marker expression and frequency of identified main populations. **c:** Visualization of log odds ratios and p values for changes in all cell-cell type interactions between experimental conditions. **d:** 1 mio EMT6-HER2 cells were injected in WT mice (i.m.). Starting from day 7 (tumor size 30–70 $mm^3$), mice were treated with $1.5 \times 10^8$ PFU of HER2-targeted and shielded adenoviral vectors (peritumorally) encoding for IL-12 on day 7, 9, 11 and 14. Lymphocyte trafficking was inhibited using FTY720 as indicated (orange arrow and line). Tumor volume on day 23 post tumor inoculation and Kaplan-Meier survival curves are shown. **e:** EMT6-HER2-engrafted mice were treated with AdV5-IL12 and AdV5-CCL5. Starting one day prior adenoviral therapy, NK cells were depleted using anti-AsialoGM1 antibody. Tumor volume on day 23 post tumor inoculation is shown. *p < 0.05, **p < 0.01, ***p < 0.001, ****p < 0.0001. Error bar values represent SEM. For survival analysis, p values were computed using the Log Rank test. For comparisons between three or more groups, one-way ANOVA with multiple comparisons was used.

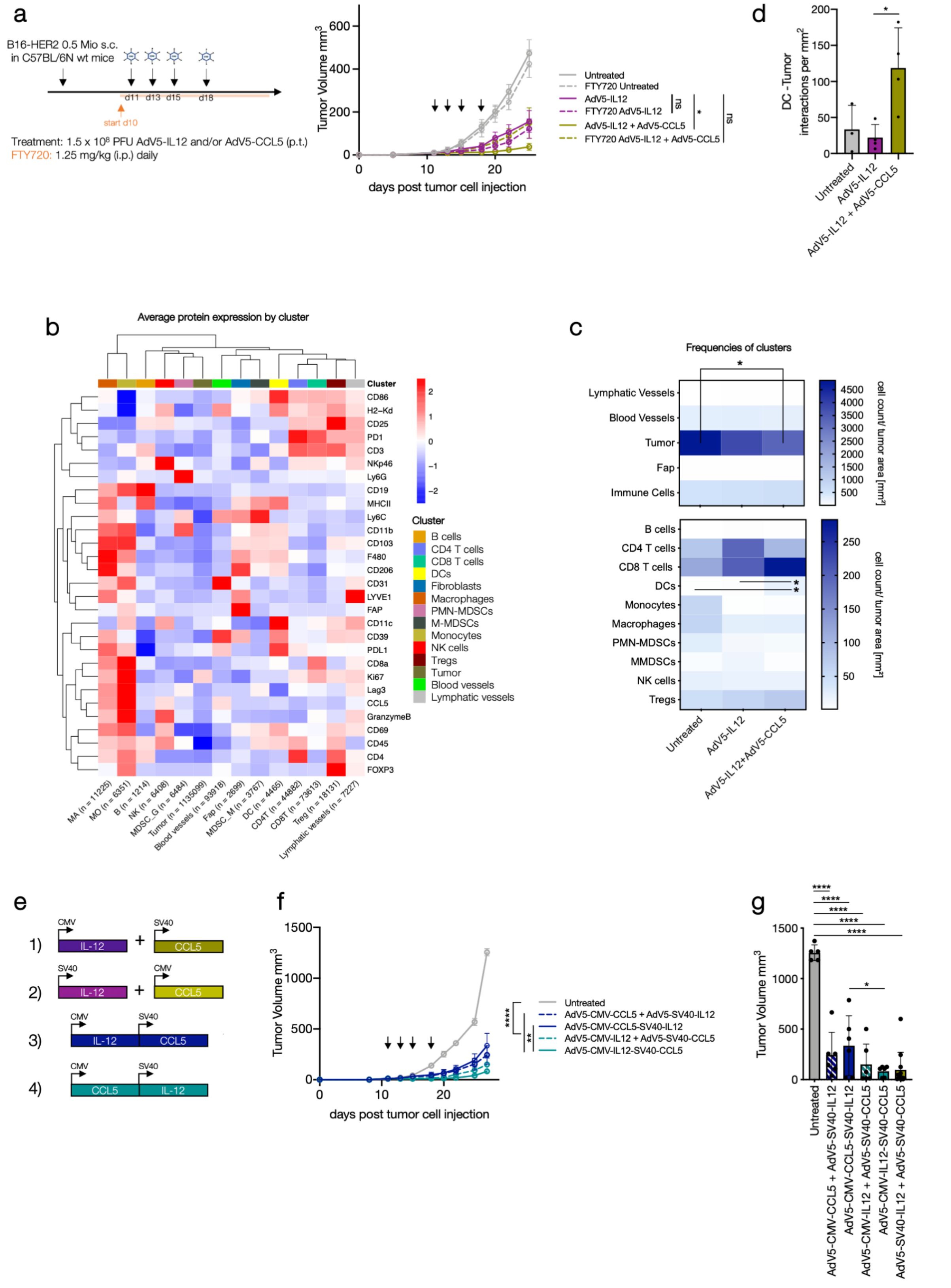

**Supplementary Fig. S5: Related to Figure 3**

**a:** Mice were treated with AdV5-IL12 and AdV5-CCL5 on day 11, 13, 15 and 18 (tumor size 30–70 mm$^3$) after B16-HER2 inoculation. Lymphocyte trafficking was inhibited using FTY720 as indicated (orange arrow and line). Tumor growth curves are shown. n = 5-6 mice per condition. Black arrows denote days of treatment. **b-d:** Wildtype (WT) mice were engrafted with 0.5 Mio B16-HER2 (s.c.). Starting from day 11 (tumor size 30–70 $mm^3$), mice were treated with $1.5x10^8$ PFU of HER2-targeted and shielded adenoviral vectors (p.t.) encoding for IL-12 and CCL5 on day 11, 13 and 15. On day 16 post inoculation, tumors were isolated, embedded in OCT and analyzed by multiparameter immuno-fluorescence microscopy. **b-c:** Heatmap showing normalized marker expression and frequency of identified main populations. **d:** Interaction count per $mm^2$ of tumor cells in close proximity to DCs. **e:** Design of combinatorial vectors compared to mixture of single viruses. **f-g:** Mice were treated with each $1.5x10^8$ PFU AdV5-IL12 and AdV5-CCL5 or $1.5x10^8$ PFU combinatorial vectors on day 11, 13, 15 and 18 (tumor size 30–70 $mm^3$) after B16-HER2 inoculation as indicated (black arrows). **f:** Tumor growth curves are shown. **g:** Quantification of tumor volume on day 27 post tumor inoculation. **f-g:** n = 5-6 mice per condition. *p < 0.05, **p < 0.01, ***p < 0.001, ****p < 0.0001. Error bar values represent SD or SEM (tumor growth curves). For comparisons between three or more groups, one-way ANOVA with multiple comparisons was used. Two-way ANOVA was used to compare tumor growth curves.

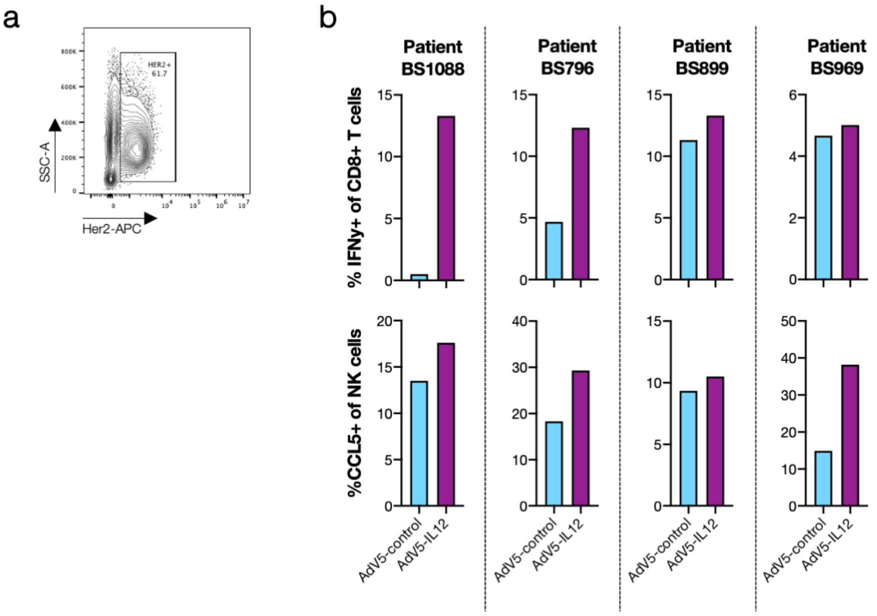


**Supplementary Fig. S6: Related to Figure 4**

HER2+ ovarian cancer samples were dissected into tumor fragments and cultivated embedded in matrigel. Tumor fragments were treated with HER2-targeted AdV5 encoding human IL-12 for 48 h (8-12 fragments per condition). **a:** Representative dot plot of HER2 expression analyzed by flow cytometry. **b:** After treatment tumor fragments per condition were pooled and analyzed by flow cytometry. Quantification of IFNγ+ CD8 T cells (CD3+, CD56-) and NK cells (CD56+, CD3-) after treatment.

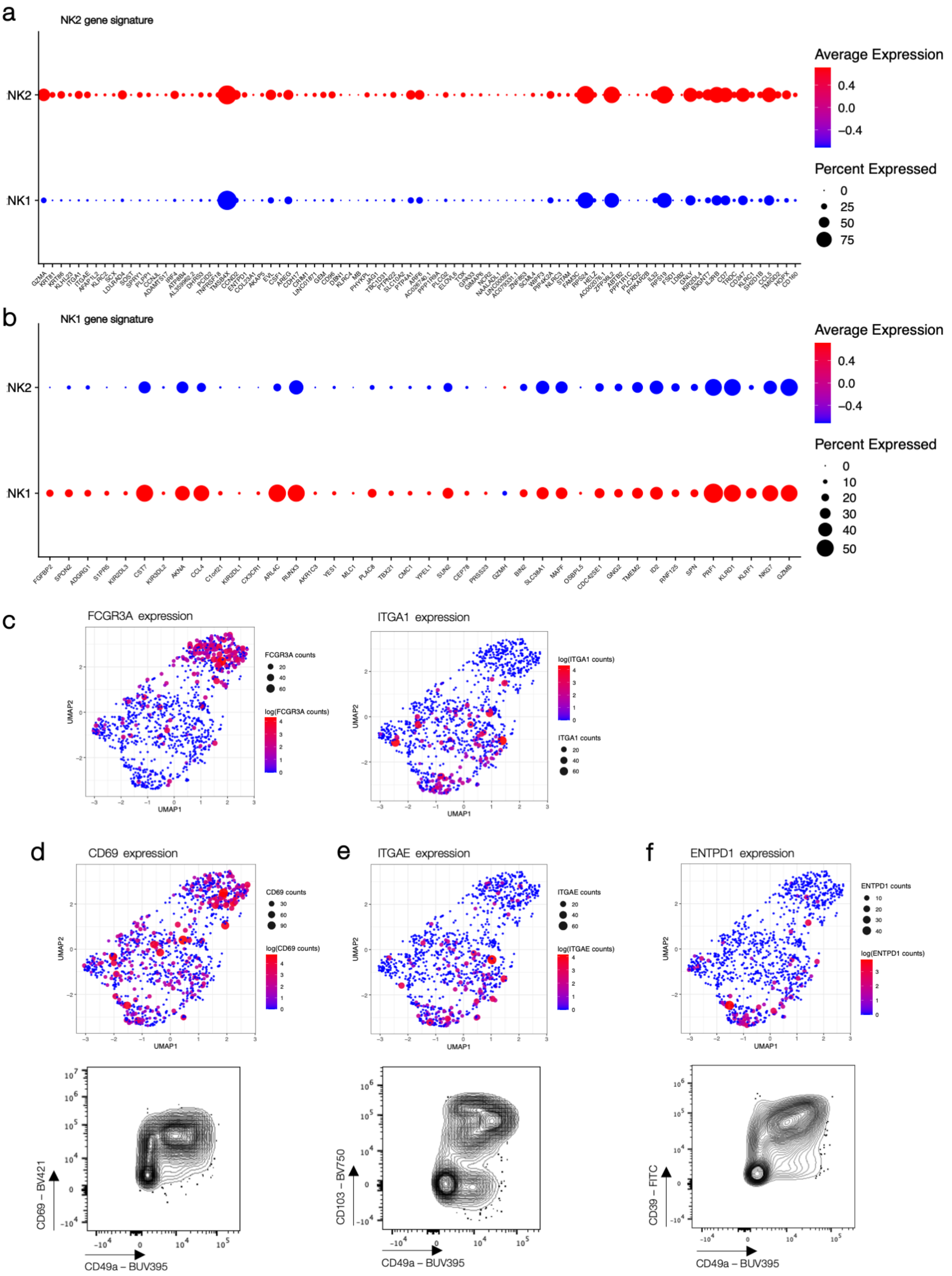


**Supplementary Fig. S7: Related to Figure 4**

**a-b:** Dotplot to visualize expression of genes in tumor-infiltrating NK cells of NSCLC patients defining generated NK2 and NK1 signature is shown. **c:** *FCGR3A* and *ITGA1* expression is visualized on UMAP projection of tumor-infiltrating NK cells of NSCLC patients. **d-e:** Comparison of gene expression of scRNASeq data set of tumor-infiltrating NK cells to protein expression analyzed by flow cytometry of NSCLC patients (CD69, *IGTAE*/CD103, *ENTPD1*/CD39).

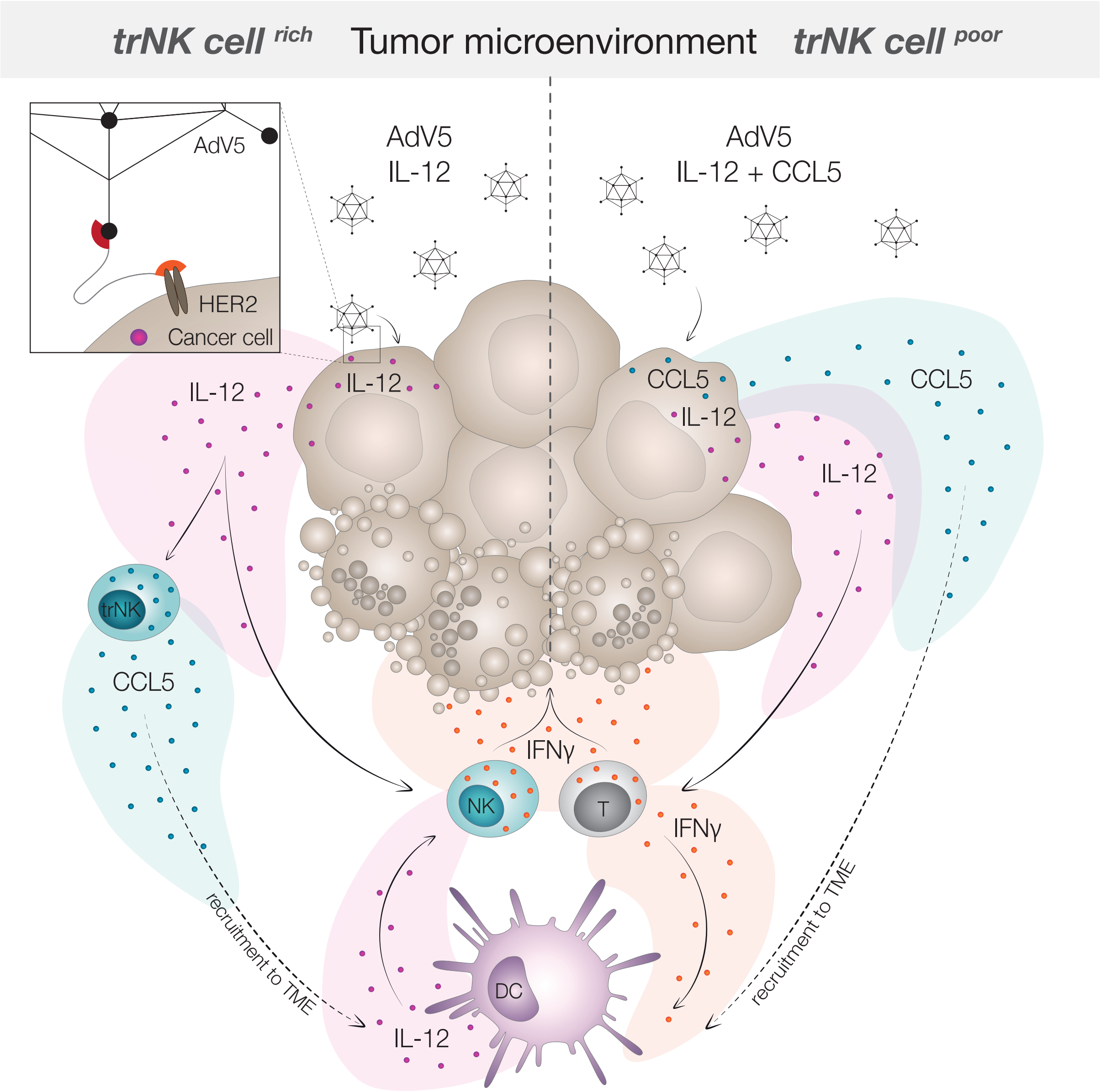


**Supplementary Fig. S8: Graphical abstract**
